# An ultra-potent synthetic nanobody neutralizes SARS-CoV-2 by locking Spike into an inactive conformation

**DOI:** 10.1101/2020.08.08.238469

**Authors:** Michael Schoof, Bryan Faust, Reuben A. Saunders, Smriti Sangwan, Veronica Rezelj, Nick Hoppe, Morgane Boone, Christian B. Billesbølle, Cristina Puchades, Caleigh M. Azumaya, Huong T. Kratochvil, Marcell Zimanyi, Ishan Deshpande, Jiahao Liang, Sasha Dickinson, Henry C. Nguyen, Cynthia M. Chio, Gregory E. Merz, Michael C. Thompson, Devan Diwanji, Kaitlin Schaefer, Aditya A. Anand, Niv Dobzinski, Beth Shoshana Zha, Camille R. Simoneau, Kristoffer Leon, Kris M. White, Un Seng Chio, Meghna Gupta, Mingliang Jin, Fei Li, Yanxin Liu, Kaihua Zhang, David Bulkley, Ming Sun, Amber M. Smith, Alexandrea N. Rizo, Frank Moss, Axel F. Brilot, Sergei Pourmal, Raphael Trenker, Thomas Pospiech, Sayan Gupta, Benjamin Barsi-Rhyne, Vladislav Belyy, Andrew W. Barile-Hill, Silke Nock, Yuwei Liu, Nevan J. Krogan, Corie Y. Ralston, Danielle L. Swaney, Adolfo García-Sastre, Melanie Ott, Marco Vignuzzi, QCRG Structural Biology Consortium, Peter Walter, Aashish Manglik

## Abstract

Without an effective prophylactic solution, infections from SARS-CoV-2 continue to rise worldwide with devastating health and economic costs. SARS-CoV-2 gains entry into host cells via an interaction between its Spike protein and the host cell receptor angiotensin converting enzyme 2 (ACE2). Disruption of this interaction confers potent neutralization of viral entry, providing an avenue for vaccine design and for therapeutic antibodies. Here, we develop single-domain antibodies (nanobodies) that potently disrupt the interaction between the SARS-CoV-2 Spike and ACE2. By screening a yeast surface-displayed library of synthetic nanobody sequences, we identified a panel of nanobodies that bind to multiple epitopes on Spike and block ACE2 interaction via two distinct mechanisms. Cryogenic electron microscopy (cryo-EM) revealed that one exceptionally stable nanobody, Nb6, binds Spike in a fully inactive conformation with its receptor binding domains (RBDs) locked into their inaccessible down-state, incapable of binding ACE2. Affinity maturation and structure-guided design of multivalency yielded a trivalent nanobody, mNb6-tri, with femtomolar affinity for SARS-CoV-2 Spike and picomolar neutralization of SARS-CoV-2 infection. mNb6-tri retains stability and function after aerosolization, lyophilization, and heat treatment. These properties may enable aerosol-mediated delivery of this potent neutralizer directly to the airway epithelia, promising to yield a widely deployable, patient-friendly prophylactic and/or early infection therapeutic agent to stem the worst pandemic in a century.

## MAIN TEXT

Over the last two decades, three zoonotic β-coronaviruses have entered the human population, causing severe respiratory symptoms with high mortality (*1-3*). The ongoing COVID-19 pandemic is caused by SARS-CoV-2, the most readily transmissible of these three coronaviruses (*4-7*). SARS-CoV-2 has wrecked the world’s economy and societies to an unprecedented extent, to date (Aug. 14, 2020) causing 751,154 reported deaths around the globe (*8*). Although public health measures have slowed its spread in many regions, infection hotspots keep reemerging. No successful vaccine or preventive treatment has yet been manufactured for any coronavirus, and the time to develop an effective and broadly available vaccine for SARS-CoV-2 remains uncertain. The development of novel therapeutic and prophylactic approaches thus remains essential, both as temporary stopgaps until an effective vaccine is generated and as permanent solutions for those segments of the population for which vaccination proves ineffective or contraindicated.

Coronavirus virions are bounded by a membrane envelope that contains ∼25 copies of the homotrimeric transmembrane spike glycoprotein (Spike) responsible for virus entry into the host cell (*9*). The surface-exposed portion of Spike is composed of two domains, S_1_ and S_2_ (*10*). The S_1_ domain mediates the interaction between virus and its host cell receptor, the angiotensin converting enzyme 2 (ACE2), while the S_2_ domain catalyzes fusion of the viral and host cell membranes (*3, 11-13*). During its biogenesis, the Spike protein is proteolytically cleaved between the S_1_ and S_2_ domains, which primes the virus for cellular entry (*10*). Contained within S_1_ is the receptor binding domain (RBD), which directly binds to ACE2. The RBD is attached to the body of Spike by a flexible region and can exist in an inaccessible down-state or an accessible up-state (*14, 15*). Binding to ACE2 requires the RBD in the up-state and enables cleavage by host proteases TMPRSS2 or cathepsin, triggering a dramatic conformational change in S_2_ that enables viral entry (*16*). In SARS-CoV-2 virions, Spike oscillates between an active, open conformation with at least one RBD in the up-state and an inactive, closed conformation with all RBDs in the down-state (*9, 11, 14, 15*).

By screening a high-complexity yeast surface-displayed library of synthetic nanobodies, we have uncovered a collection of nanobodies that block the Spike-ACE2 interaction. Biochemical and structural studies revealed that two classes of these nanobodies act in distinct ways to prevent ACE2 binding. Combining affinity maturation and structure-guided multimerization, we optimized these agents and generated Spike binders that match or exceed the potency of most monoclonal antibodies disclosed to date. Our lead neutralizing molecule, mNb6-tri, blocks SARS-CoV-2 entry in human cells at picomolar efficacy and withstands aerosolization, lyophilization, and elevated temperatures. mNb6-tri provides a promising approach to deliver a potent SARS-CoV-2 neutralizing molecule directly to the airways for prophylaxis or therapy.

## RESULTS

### Synthetic nanobodies that disrupt Spike-ACE2 interaction

To isolate nanobodies that neutralize SARS-CoV-2, we screened a yeast surface-displayed library of >2×10^9^ synthetic nanobody sequences. Our strategy was to screen for binders to the full Spike protein ectodomain, in order to capture not only those nanobodies that would compete by binding to the ACE2-binding site on the RBD directly but also those that might bind elsewhere on Spike and block ACE2 interaction through indirect mechanisms. We used a mutant form of SARS-CoV-2 Spike (Spike*,) as the antigen (*15*). Spike* lacks one of the two activating proteolytic cleavage sites between the S_1_ and S_2_ domains and introduces two mutations to stabilize the pre-fusion conformation. Spike* expressed in mammalian cells binds ACE2 with a K_D_ = 44 nM (Supplementary Fig. 1), consistent with previous reports (*17*). Next, we labeled Spike* with biotin or with fluorescent dyes and selected nanobody-displaying yeast over multiple rounds, first by magnetic bead binding and then by fluorescence-activated cell sorting (Fig. 1A).

**Figure 1.**
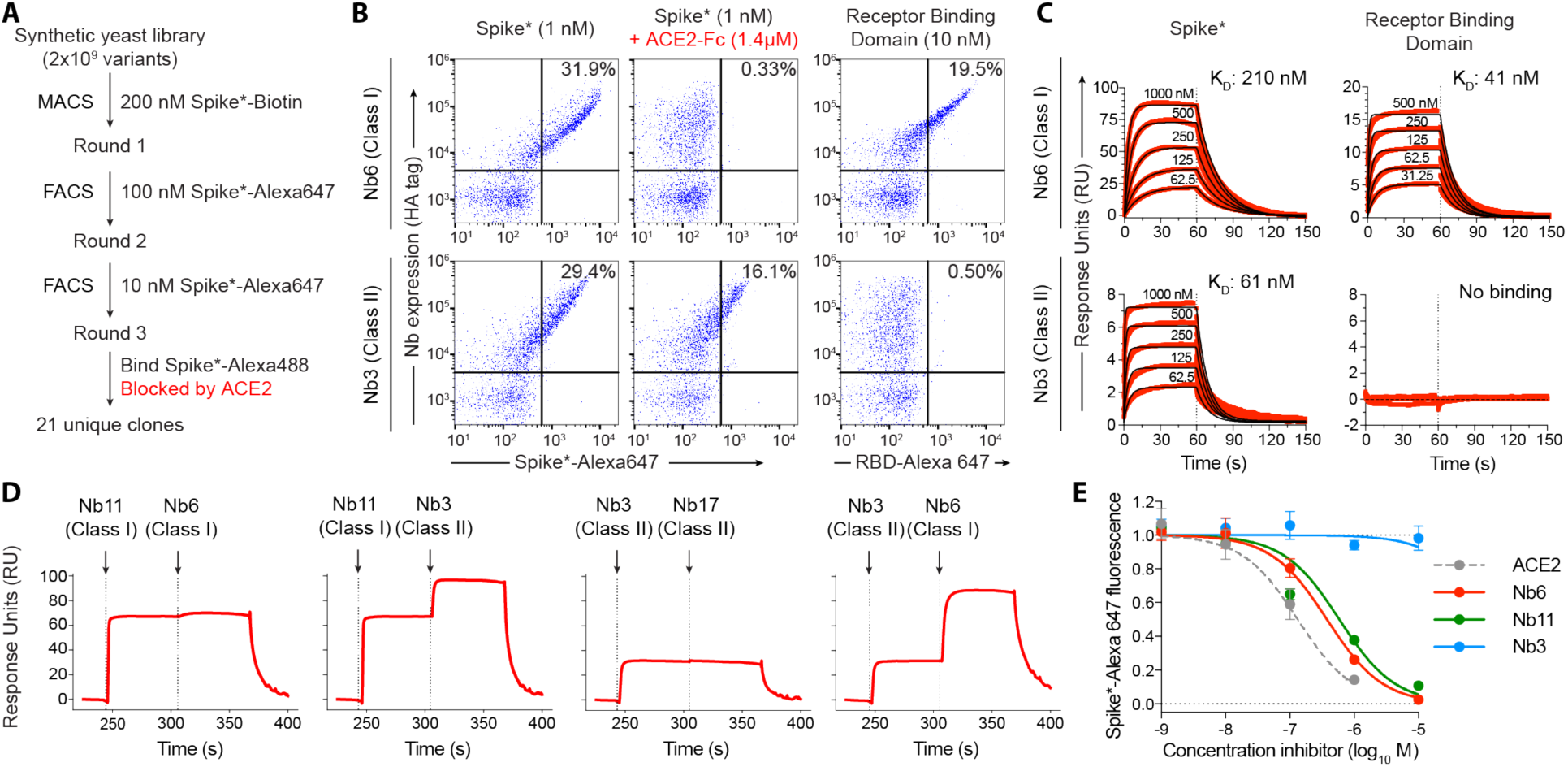
Discovery of two distinct classes of anti-Spike nanobodies. **A**, Selection strategy for identification of anti-Spike nanobodies that disrupt Spike-ACE2 interactions using magnetic bead selections (MACS) or fluorescence activated cell sorting (FACS). **B**, Flow cytometry of yeast displaying Nb6 (a Class I nanobody) or Nb3 (a Class II nanobody). Nb6 binds Spike*-Alexa 647 and receptor binding domain (RBD-Alexa 647). Nb6 binding to Spike* is completely disrupted by an excess (1.4 µM) of ACE2-Fc. Nb3 binds Spike*, but not the RBD. Nb3 binding to Spike* is partially decreased by ACE2-Fc. **C**, SPR of Nb6 and Nb3 binding to either Spike* or RBD. Red traces are raw data and global kinetic fits are shown in black. Nb3 shows no binding to RBD. **D**, SPR experiments with immobilized Spike* show that Class I and Class II nanobodies can bind Spike* simultaneously. By contrast, two Class I nanobodies or Class II nanobodies do not bind simultaneously. **E**, Nanobody inhibition of 1 nM Spike*-Alexa 647 binding to ACE2 expressing HEK293T cells. n = 3 (ACE2, Nb3) or 5 (Nb6, Nb11) biological replicates. All error bars represent s.e.m.

Three rounds of selection yielded 21 unique nanobodies that bound Spike* and showed decreased Spike* binding in the presence of ACE2. Closer inspection of their binding properties revealed that these nanobodies fall into two distinct classes. One group (Class I) binds the RBD and competes with ACE2 (Fig. 1B). A prototypical example of this class is nanobody Nb6, which binds to Spike* and to RBD alone with a K_D_ of 210 nM and 41 nM, respectively (Fig. 1C; Table 1). Another group (Class II), exemplified by nanobody Nb3, binds to Spike* (K_D_ = 61 nM), but displays no binding to RBD alone (Fig. 1C, Table 1). In the presence of excess ACE2, binding of Nb6 and other Class I nanobodies is blocked entirely, whereas binding of Nb3 and other Class II nanobodies is decreased only moderately (Fig. 1B). These results suggest that Class I nanobodies target the RBD to block ACE2 binding, whereas Class II nanobodies target other epitopes and decrease ACE2 interaction with Spike allosterically or through steric interference. Indeed, surface plasmon resonance (SPR) experiments demonstrate that Class I and Class II nanobodies can bind Spike* simultaneously (Fig. 1D).

**Table 1.**
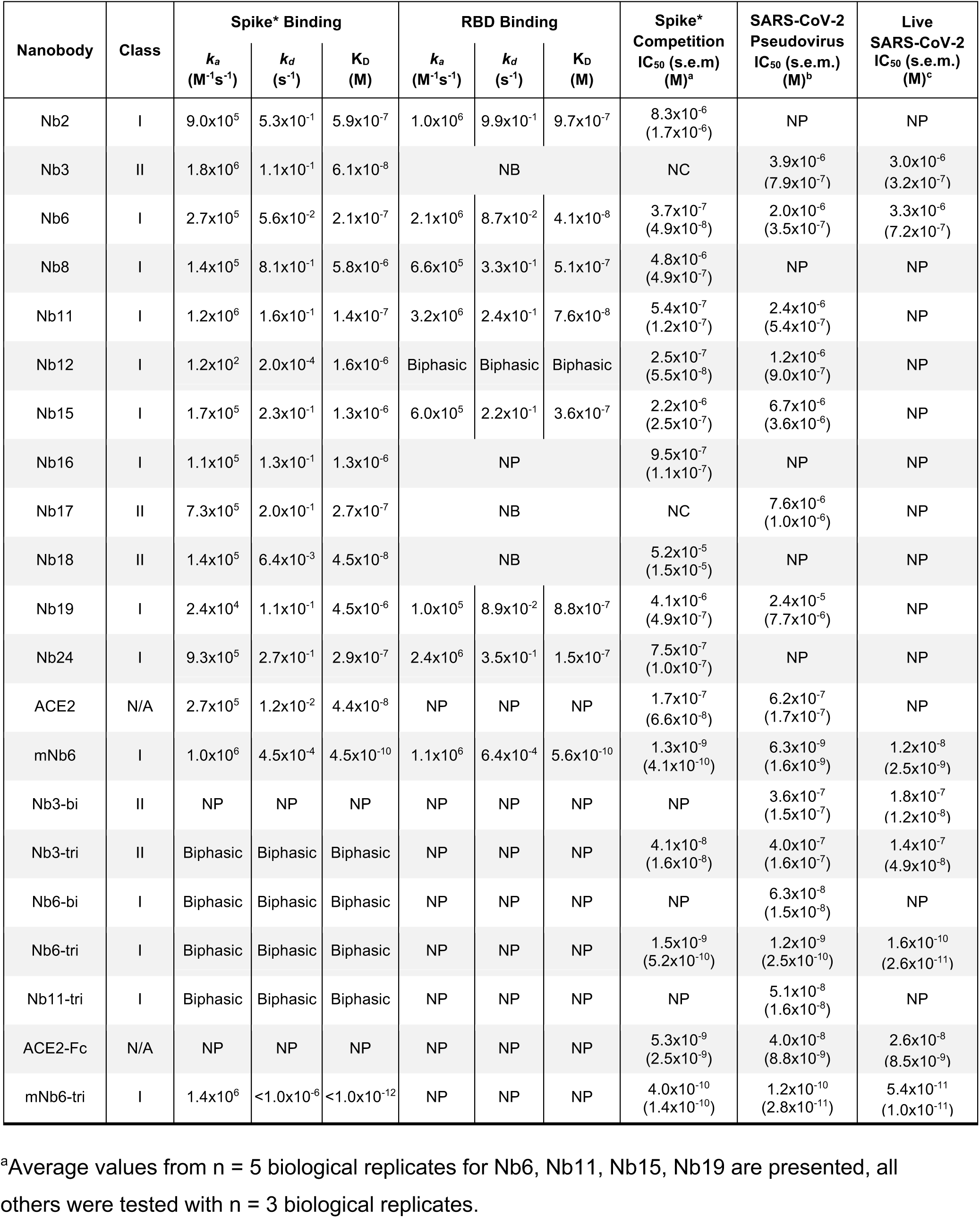
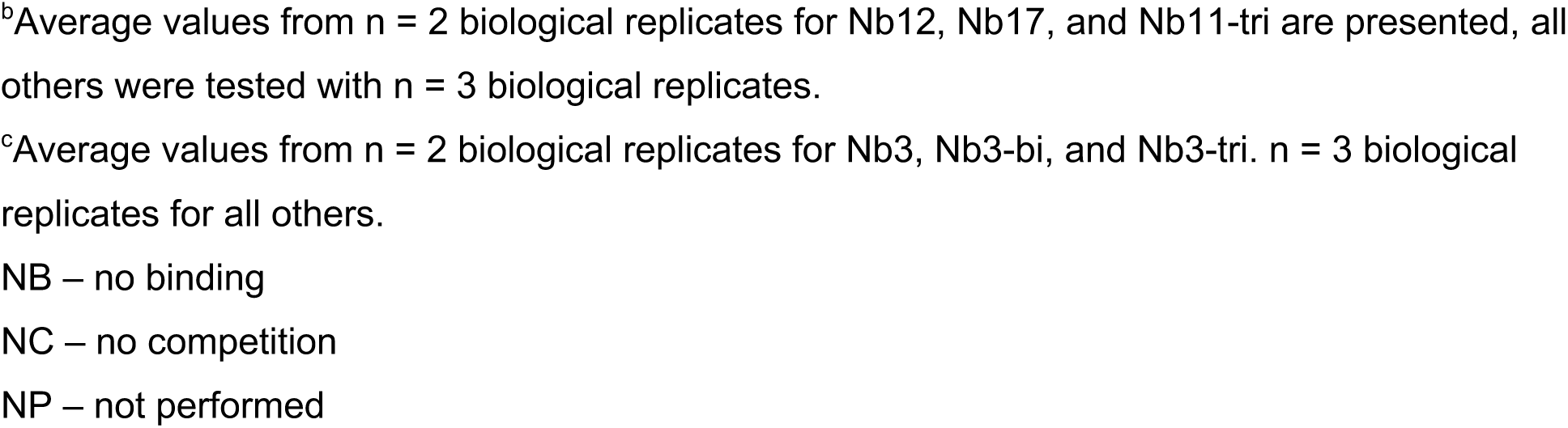
Anti-Spike nanobody affinity and neutralization potency.

| Nanobody | Class | Spike* Binding |  |  | RBD Binding |  |  | Spike* Competition<br>IC<sub>50</sub> (s.e.m)<br>(M)<sup>a</sup> | SARS-CoV-2 Pseudovirus<br>IC<sub>50</sub> (s.e.m.)<br>(M)<sup>b</sup> | Live SARS-CoV-2<br>IC<sub>50</sub> (s.e.m.)<br>(M)<sup>c</sup> |
| --- | --- | --- | --- | --- | --- | --- | --- | --- | --- | --- |
|  |  | *k<sub>a</sub>*<br>(M<sup>-1</sup>s<sup>-1</sup>) | *k<sub>d</sub>*<br>(s<sup>-1</sup>) | *K<sub>D</sub>*<br>(M) | *k<sub>a</sub>*<br>(M<sup>-1</sup>s<sup>-1</sup>) | *k<sub>d</sub>*<br>(s<sup>-1</sup>) | *K<sub>D</sub>*<br>(M) |  |  |  |
| Nb2 | I | 9.0x10<sup>5</sup> | 5.3x10<sup>-1</sup> | 5.9x10<sup>-7</sup> | 1.0x10<sup>6</sup> | 9.9x10<sup>-1</sup> | 9.7x10<sup>-7</sup> | 8.3x10<sup>-6</sup><br>(1.7x10<sup>-6</sup>) | NP | NP |
| Nb3 | II | 1.8x10<sup>6</sup> | 1.1x10<sup>-1</sup> | 6.1x10<sup>-8</sup> | NB |  |  | NC | 3.9x10<sup>-6</sup><br>(7.9x10<sup>-7</sup>) | 3.0x10<sup>-6</sup><br>(3.2x10<sup>-7</sup>) |
| Nb6 | I | 2.7x10<sup>5</sup> | 5.6x10<sup>-2</sup> | 2.1x10<sup>-7</sup> | 2.1x10<sup>6</sup> | 8.7x10<sup>-2</sup> | 4.1x10<sup>-8</sup> | 3.7x10<sup>-7</sup><br>(4.9x10<sup>-8</sup>) | 2.0x10<sup>-6</sup><br>(3.5x10<sup>-7</sup>) | 3.3x10<sup>-6</sup><br>(7.2x10<sup>-7</sup>) |
| Nb8 | I | 1.4x10<sup>5</sup> | 8.1x10<sup>-1</sup> | 5.8x10<sup>-6</sup> | 6.6x10<sup>5</sup> | 3.3x10<sup>-1</sup> | 5.1x10<sup>-7</sup> | 4.8x10<sup>-6</sup><br>(4.9x10<sup>-7</sup>) | NP | NP |
| Nb11 | I | 1.2x10<sup>6</sup> | 1.6x10<sup>-1</sup> | 1.4x10<sup>-7</sup> | 3.2x10<sup>6</sup> | 2.4x10<sup>-1</sup> | 7.6x10<sup>-8</sup> | 5.4x10<sup>-7</sup><br>(1.2x10<sup>-7</sup>) | 2.4x10<sup>-6</sup><br>(5.4x10<sup>-7</sup>) | NP |
| Nb12 | I | 1.2x10<sup>2</sup> | 2.0x10<sup>-4</sup> | 1.6x10<sup>-6</sup> | Biphasic | Biphasic | Biphasic | 2.5x10<sup>-7</sup><br>(5.5x10<sup>-8</sup>) | 1.2x10<sup>-6</sup><br>(9.0x10<sup>-7</sup>) | NP |
| Nb15 | I | 1.7x10<sup>5</sup> | 2.3x10<sup>-1</sup> | 1.3x10<sup>-6</sup> | 6.0x10<sup>5</sup> | 2.2x10<sup>-1</sup> | 3.6x10<sup>-7</sup> | 2.2x10<sup>-6</sup><br>(2.5x10<sup>-7</sup>) | 6.7x10<sup>-6</sup><br>(3.6x10<sup>-6</sup>) | NP |
| Nb16 | I | 1.1x10<sup>5</sup> | 1.3x10<sup>-1</sup> | 1.3x10<sup>-6</sup> | NP |  |  | 9.5x10<sup>-7</sup><br>(1.1x10<sup>-7</sup>) | NP | NP |
| Nb17 | II | 7.3x10<sup>5</sup> | 2.0x10<sup>-1</sup> | 2.7x10<sup>-7</sup> | NB |  |  | NC | 7.6x10<sup>-6</sup><br>(1.0x10<sup>-6</sup>) | NP |
| Nb18 | II | 1.4x10<sup>5</sup> | 6.4x10<sup>-3</sup> | 4.5x10<sup>-8</sup> | NB |  |  | 5.2x10<sup>-5</sup><br>(1.5x10<sup>-5</sup>) | NP | NP |
| Nb19 | I | 2.4x10<sup>4</sup> | 1.1x10<sup>-1</sup> | 4.5x10<sup>-6</sup> | 1.0x10<sup>5</sup> | 8.9x10<sup>-2</sup> | 8.8x10<sup>-7</sup> | 4.1x10<sup>-6</sup><br>(4.9x10<sup>-7</sup>) | 2.4x10<sup>-5</sup><br>(7.7x10<sup>-6</sup>) | NP |
| Nb24 | I | 9.3x10<sup>5</sup> | 2.7x10<sup>-1</sup> | 2.9x10<sup>-7</sup> | 2.4x10<sup>6</sup> | 3.5x10<sup>-1</sup> | 1.5x10<sup>-7</sup> | 7.5x10<sup>-7</sup><br>(1.0x10<sup>-7</sup>) | NP | NP |
| ACE2 | N/A | 2.7x10<sup>5</sup> | 1.2x10<sup>-2</sup> | 4.4x10<sup>-8</sup> | NP | NP | NP | 1.7x10<sup>-7</sup><br>(6.6x10<sup>-8</sup>) | 6.2x10<sup>-7</sup><br>(1.7x10<sup>-7</sup>) | NP |
| mNb6 | I | 1.0x10<sup>6</sup> | 4.5x10<sup>-4</sup> | 4.5x10<sup>-10</sup> | 1.1x10<sup>6</sup> | 6.4x10<sup>-4</sup> | 5.6x10<sup>-10</sup> | 1.3x10<sup>-9</sup><br>(4.1x10<sup>-10</sup>) | 6.3x10<sup>-9</sup><br>(1.6x10<sup>-9</sup>) | 1.2x10<sup>-8</sup><br>(2.5x10<sup>-9</sup>) |
| Nb3-bi | II | NP | NP | NP | NP | NP | NP | NP | 3.6x10<sup>-7</sup><br>(1.5x10<sup>-7</sup>) | 1.8x10<sup>-7</sup><br>(1.2x10<sup>-8</sup>) |
| Nb3-tri | II | Biphasic | Biphasic | Biphasic | NP | NP | NP | 4.1x10<sup>-8</sup><br>(1.6x10<sup>-8</sup>) | 4.0x10<sup>-7</sup><br>(1.6x10<sup>-7</sup>) | 1.4x10<sup>-7</sup><br>(4.9x10<sup>-8</sup>) |
| Nb6-bi | I | Biphasic | Biphasic | Biphasic | NP | NP | NP | NP | 6.3x10<sup>-8</sup><br>(1.5x10<sup>-8</sup>) | NP |
| Nb6-tri | I | Biphasic | Biphasic | Biphasic | NP | NP | NP | 1.5x10<sup>-9</sup><br>(5.2x10<sup>-10</sup>) | 1.2x10<sup>-9</sup><br>(2.5x10<sup>-10</sup>) | 1.6x10<sup>-10</sup><br>(2.6x10<sup>-11</sup>) |
| Nb11-tri | I | Biphasic | Biphasic | Biphasic | NP | NP | NP | NP | 5.1x10<sup>-8</sup><br>(1.6x10<sup>-8</sup>) | NP |
| ACE2-Fc | N/A | NP | NP | NP | NP | NP | NP | 5.3x10<sup>-9</sup><br>(2.5x10<sup>-9</sup>) | 4.0x10<sup>-8</sup><br>(8.8x10<sup>-9</sup>) | 2.6x10<sup>-8</sup><br>(8.5x10<sup>-9</sup>) |
| mNb6-tri | I | 1.4x10<sup>6</sup> | <1.0x10<sup>-6</sup> | <1.0x10<sup>-12</sup> | NP | NP | NP | 4.0x10<sup>-10</sup><br>(1.4x10<sup>-10</sup>) | 1.2x10<sup>-10</sup><br>(2.8x10<sup>-11</sup>) | 5.4x10<sup>-11</sup><br>(1.0x10<sup>-11</sup>) |

<sup>a</sup>Average values from n = 5 biological replicates for Nb6, Nb11, Nb15, Nb19 are presented, all others were tested with n = 3 biological replicates.
<sup>b</sup>Average values from n = 2 biological replicates for Nb12, Nb17, and Nb11-tri are presented, all others were tested with n = 3 biological replicates.
<sup>c</sup>Average values from n = 2 biological replicates for Nb3, Nb3-bi, and Nb3-tri. n = 3 biological replicates for all others.
NB – no binding
NC – no competition
NP – not performed

Analysis of the kinetic rate constants for Class I nanobodies revealed a consistently greater association rate constant (*k*_*a*_) for nanobody binding to the isolated RBD than to full-length Spike* (Table 1), which suggests that RBD accessibility influences the K_D_. We next tested the efficacy of our nanobodies, both Class I and Class II, to inhibit binding of fluorescently labeled Spike* to ACE2-expressing HEK293 cells (Table 1, Fig. 1E). Class I nanobodies emerged with highly variable activity in this assay with Nb6 and Nb11 as two of the most potent clones with IC_50_ values of 370 and 540 nM, respectively (Table 1). For unexplained reasons, Class II nanobodies showed little to no activity in this assay (Table 1, Fig. 1E).

Going forward, we prioritized two Class I nanobodies, Nb6 and Nb11, that combine potent Spike* binding with relatively small differences in *k*_*a*_ between binding to Spike* or RBD. We reasoned that the epitopes recognized by Nb6 and Nb11 would be more readily accessible in the Spike protein on intact virions. For Class II nanobodies we prioritized Nb3 because of its optimal stability and yield during purification.

### Nb6 and Nb11 target the RBD and directly compete with ACE2

To define the binding sites of Nb6 and Nb11, we determined their cryogenic electron microscopy (cryo-EM) structures bound to Spike* (Fig. 2A-B, Supplementary Fig. 2-4, Supplementary Table 1). Both nanobodies recognize RBD epitopes that overlap the ACE2 binding site (Fig. 2E). For Nb6 and Nb11, we resolved nanobody binding to both the open and closed conformations of Spike*. We obtained a 3.0 Å map of Nb6 bound to closed Spike*, which enabled modeling of the Nb6-Spike* complex (Fig. 2A), including the complementarity determining regions (CDRs). We also obtained lower resolution maps for Nb6 bound to open Spike* (3.8 Å), Nb11 bound to open Spike* (4.2 Å), and Nb11 bound to closed Spike* (3.7 Å). For these lower resolution maps, we could define the nanobody’s binding orientation but not accurately model the CDRs.

**Figure 2.**
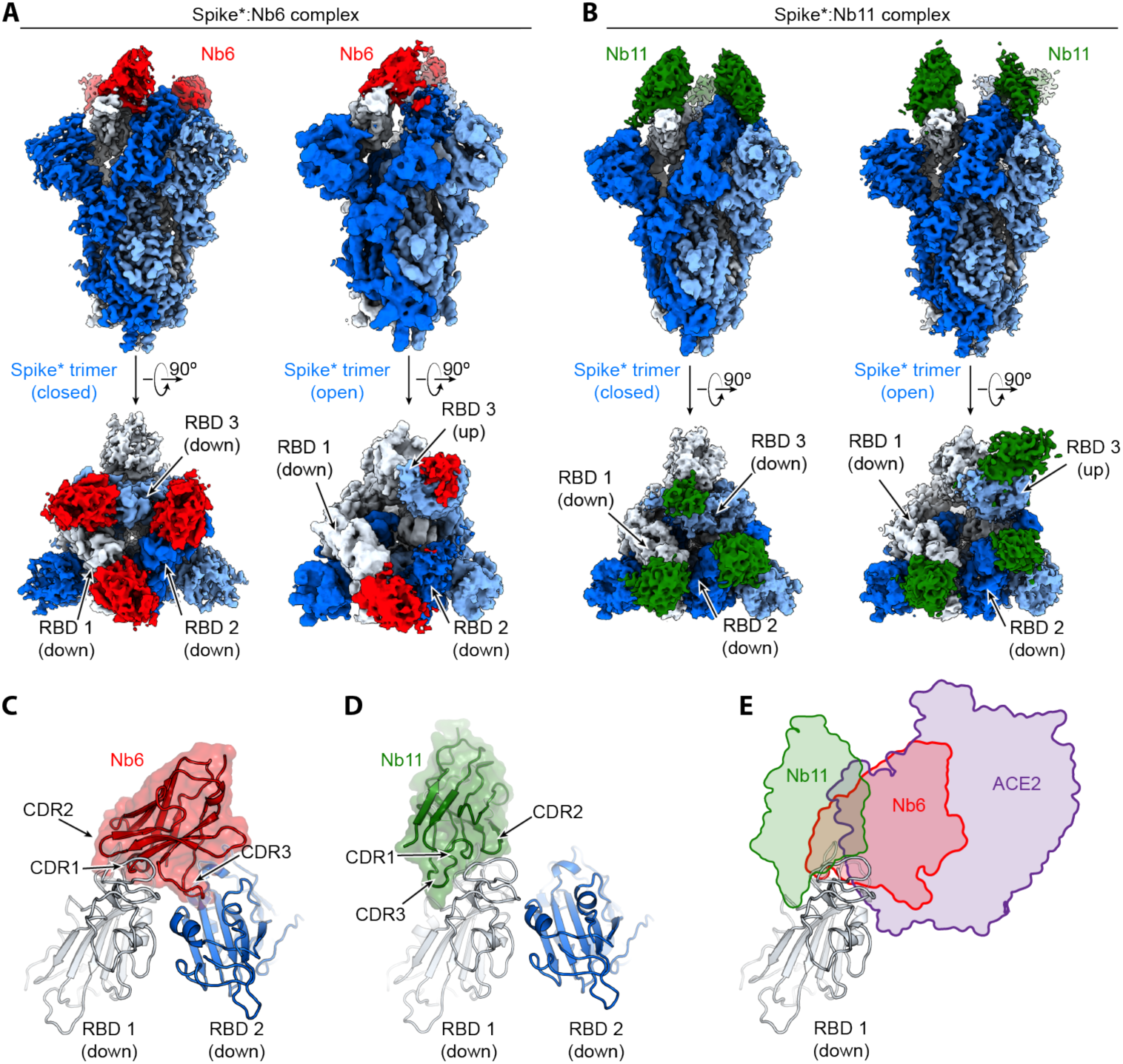
Cryo-EM structures of Nb6 and Nb11 bound to Spike. **A**, Cryo-EM maps of Spike*-Nb6 complex in either closed (left) or open (right) Spike* conformation. **B**, Cryo-EM maps of Spike*-Nb11 complex in either closed (left) or open (right) Spike* conformation. The top views show receptor binding domain (RBD) up-or down-states. **C**, Nb6 straddles the interface of two down-state RBDs, with CDR3 reaching over to an adjacent RBD. **D**, Nb11 binds a single RBD in the down-state (displayed) or similarly in the up-state. No cross-RBD contacts are made by Nb11 in either RBD up-or down-state. **E**, Comparison of RBD epitopes engaged by ACE2 (purple), Nb6 (red), or Nb11 (green). Both Nb11 and Nb6 directly compete with ACE2 binding.

Nb6 bound to closed Spike* straddles the interface between two adjacent RBDs. The majority of the contacting surfaces are contributed by CDR1 and CDR2 of Nb6 (Fig. 2C). CDR3 contacts the adjacent RBD that is counterclockwise positioned when viewed from the top of Spike* (Fig. 2C). The binding of one Nb6 therefore stabilizes two adjacent RBDs in the down-state. We surmise that this initial binding event pre-organizes the binding site for a second and third Nb6 molecule to stabilize the closed Spike* conformation. Indeed, binding of two Nb6 molecules would lock all three RBDs into the down-state, thus highly favoring binding of a third Nb6 because binding would not entail any further entropic cost. By contrast, Nb11 bound to down-state RBDs only contacts a single RBD (Fig. 2D).

### Nb3 interacts with the Spike S_1_ domain external to the RBD

Our attempts to determine the binding site of Nb3 by cryo-EM proved unsuccessful. We therefore turned to radiolytic hydroxyl radical footprinting to determine potential binding sites for Nb3. Spike*, either apo or bound to Nb3, was exposed to 5-50 milliseconds of synchrotron X-ray radiation to label solvent-exposed amino acids with hydroxyl radicals. Radical-labeled amino acids were subsequently identified and quantified by mass spectrometry of trypsin/Lys-C or Glu-C protease digested Spike*(*18*). Two neighboring surface residues on the S_1_ domain of Spike (M177 and H207) emerged as highly protected sites in the presence of Nb3 (Supplementary Fig. 5). The degree of protection is consistent with prior observations of antibody-antigen interactions by hydroxyl radical footprinting (*19*). Both M177 and H207 are greater than 40 Å distant from the ACE2 binding site on the RBD, suggesting that Nb3 may inhibit Spike-ACE2 interactions through allosteric means.

### Rationally engineered multivalency increases potency

The structure of Nb6 bound to closed Spike* enabled us to engineer bivalent and trivalent nanobodies predicted to lock all RBDs in the down-state. To this end, we inserted flexible Gly-Ser linkers of either 15 or 20 amino acids to span the 52 Å distance between adjacent Nb6 monomers bound to down-state RBDs in closed Spike* (Supplementary Fig. 6). Both linker lengths are too short to span the distance (72 Å) between Nb6 bound to a down-state RBD and an up-state RBD that would co-exist in an open Spike. Moreover, binding of three RBDs in the previously reported conformation of Nb6-bound open Spike* would be physically impossible even with longer linker length because of steric clashes (Supplementary Fig. 6). By contrast, the minimum distance between adjacent Nb11 monomers bound to either open or closed Spike* is 68 Å (Supplementary Fig. 6). We therefore predicted that multivalent binding by Nb6 constructs would display significantly slowed dissociation rates due to the enhanced avidity afforded by Spike’s trimeric architecture.

We assessed multivalent Nb6 binding to Spike* by SPR. Both bivalent Nb6 with a 15 amino acid linker (Nb6-bi) and trivalent Nb6 with two 20 amino acid linkers (Nb6-tri) dissociate from Spike* in a biphasic manner. The dissociation phase can be fitted to two components: a fast phase with kinetic rate constants *k*_*d1*_ of 2.7×10^−2^ s^-1^ for Nb6-bi and 2.9×10^−2^ s^-1^ for Nb6-tri, which are of the same magnitude as that observed for monovalent Nb6 (*k*_*d*_ = 5.6×10^−2^ s^-1^) and a slow phase that is dependent on avidity (*k*_*d2*_ = 3.1×10^−4^ for Nb6-bi and *k*_*d2*_ < 1.0×10^−6^ s^-1^ for Nb6-tri, respectively) (Fig. 3A). The relatively similar *k*_*d*_ for the fast phase suggests that a fraction of the observed binding for the multivalent constructs is nanobody binding to a single Spike* RBD. By contrast, the slow dissociation phase of Nb6-bi and Nb6-tri indicates engagement of two or three RBDs. We observed no dissociation for the slow phase of Nb6-tri over 10 minutes, indicating an upper boundary for *k*_*d2*_ of 1×10^−6^ s^-1^ and subpicomolar affinity. This measurement remains an upper-bound estimate rather than an accurate measurement because the technique is limited by the intrinsic dissociation rate of Spike* from the chip imposed by the chemistry used to immobilize Spike*.

**Figure 3.**
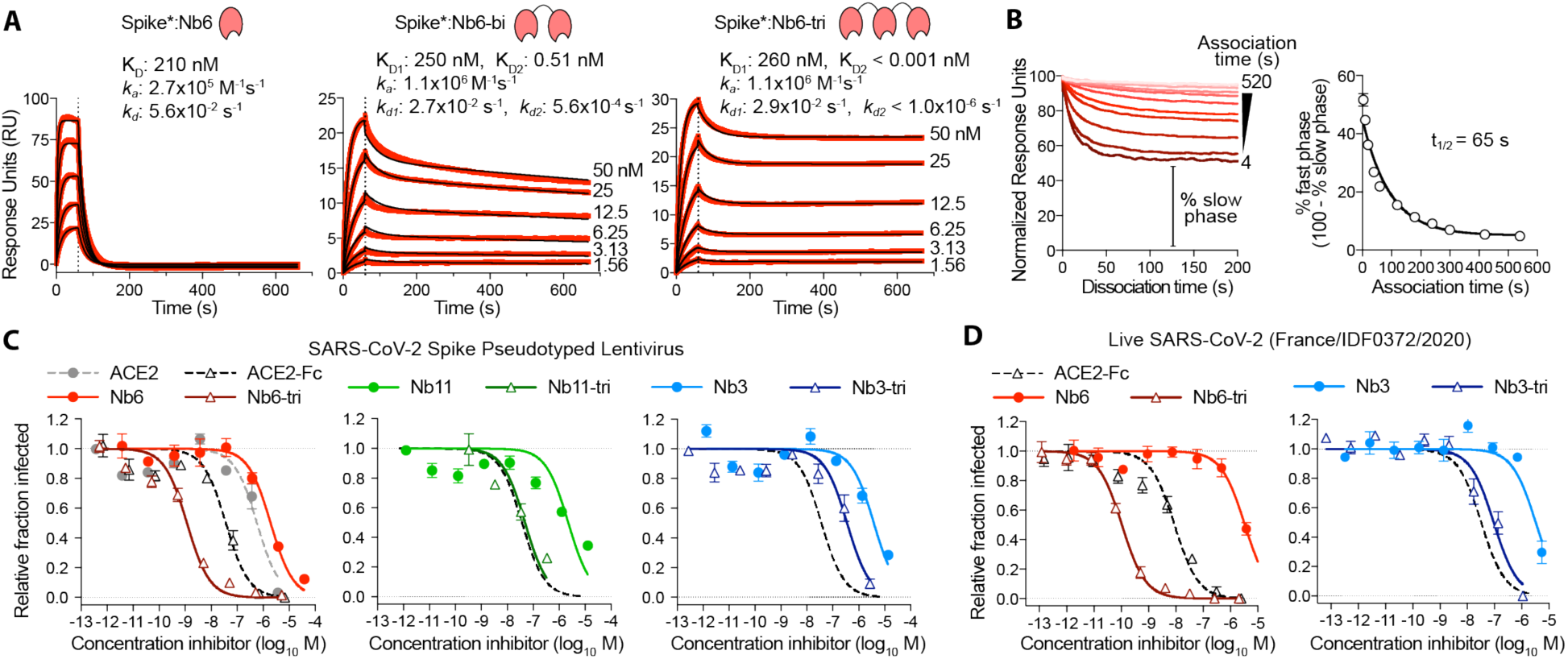
Multivalency improves nanobody affinity and inhibitory efficacy. **A**, SPR of Nb6 and multivalent variants. Red traces show raw data and black lines show global kinetic fit for Nb6 and independent fits for association and dissociation phases for Nb6-bi and Nb6-tri. **B**, Dissociation phase SPR traces for Nb6-tri after variable association time ranging from 4 to 520 s. Curves were normalized to maximal signal at the beginning of the dissociation phase. Percent fast phase is plotted as a function of association time (right) with a single exponential fit. n = 3 independent biological replicates. **C**, Inhibition of pseudotyped lentivirus infection of ACE2 expressing HEK293T cells. n = 3 biological replicates for all but Nb11-tri (n = 2) **D**, Inhibition of live SARS-CoV-2 virus. Representative biological replicate with n = 3 (right panel) or 4 (left panel) technical replicates per concentration. n = 3 biological replicates for all but Nb3 and Nb3-tri (n = 2). All error bars represent s.e.m.

We reasoned that the biphasic dissociation behavior could be explained by a slow interconversion between up- and down-state RBDs, with conversion to the more stable down-state required for full trivalent binding. According to this view, a single domain of Nb6-tri engaged with an up-state RBD would dissociate rapidly. The system would then re-equilibrate as the RBD flips into the down-state, eventually allowing Nb6-tri to trap all RBDs in closed Spike*. To test this notion directly, we varied the time allowed for Nb6-tri binding to Spike*. Indeed, we observed an exponential decrease in the percent fast-phase with a t_1/2_ of 65 s (Fig. 3B), which, we surmise, reflects the timescale of conversion between the RBD up- and down-states in Spike*. Taken together, dimerization and trimerization of Nb6 afforded 750-fold and >200,000-fold gains in K_D_, respectively.

### Class I and II nanobodies prevent SARS-CoV-2 infection

We next tested the neutralization activity of trivalent versions of our top Class I (Nb6 and Nb11) and Class II (Nb3) nanobodies against SARS-CoV-2 pseudotyped lentivirus. In this assay, SARS-CoV-2 Spike is expressed as a surface protein on a lentiviral particle that contains a ZsGreen reporter gene, which is integrated and expressed upon successful viral entry into cells harboring the ACE2 receptor (*20*). Nb6 and Nb11 inhibited pseudovirus infection with IC_50_ values of 2.0 µM and 2.4 µM, respectively, and Nb3 inhibited pseudovirus infection with an IC_50_ of 3.9 µM (Fig. 3C, Table 1). Nb6-tri shows a 2000-fold enhancement of inhibitory activity, with an IC_50_ of 1.2 nM, whereas trimerization of Nb11 and Nb3 resulted in more modest gains of 40- and 10-fold (51 nM and 400 nM), respectively (Fig. 3C).

We next confirmed these neutralization activities with a viral plaque assay using live SARS-CoV-2 virus infection of VeroE6 cells. Consistent with its activity against pseudotyped lentivirus, Nb6-tri proved exceptionally potent, neutralizing SARS-CoV-2 with an average IC_50_ of 160 pM (Fig. 3D). Nb3-tri neutralized SARS-CoV-2 with an average IC_50_ of 140 nM (Fig. 3D).

### Affinity maturation yields a femtomolar K_D_ Spike inhibitor

We further optimized the potency of Nb6 by selecting high-affinity variants. To this end, we prepared a new library, starting with the Nb6 coding sequence, in which we varied each amino acid position of all three CDRs by saturation mutagenesis (Fig. 4A). After two rounds of magnetic bead-based selection, we isolated a population of high-affinity clones. Sequencing revealed two highly penetrant mutations: I27Y in CDR1 and P105Y in CDR3. We incorporated these two mutations into Nb6 to generate matured Nb6 (mNb6), which binds with 500-fold increased affinity to Spike* as measured by SPR (Fig. 4B). As a monomer, mNb6 inhibits both pseudovirus and live SARS-CoV-2 infection with low nanomolar potency, a ∼200-fold improvement compared to Nb6 (Fig. 4I-J, Table 1).

**Figure 4.**
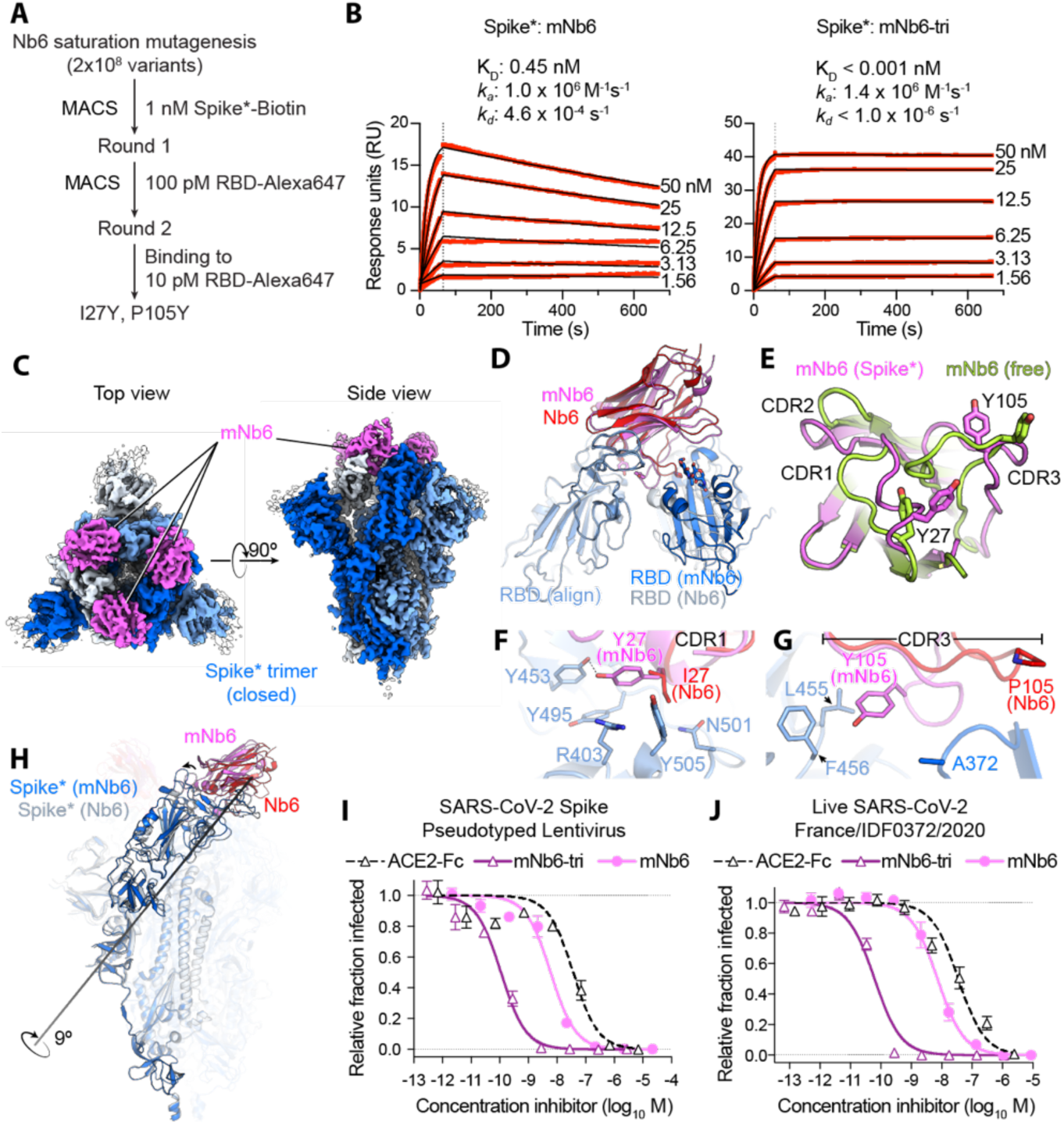
Affinity maturation of Nb6 yields a picomolar SARS-CoV-2 neutralizing molecule. **A**, A saturation mutagenesis library of Nb6 was subjected to two rounds of selection to identify consensus mutations I27Y and P105Y. **B**, SPR of mNb6 and mNb6-tri binding to immobilized Spike*. Red traces show raw data and black lines show global kinetic fit. No dissociation was observed for mNb6-tri over 10 minutes. **C**, Cryo-EM structure of Spike*-mNb6 complex. **D**, Comparison of receptor binding domain (RBD) engagement by Nb6 and mNb6. One RBD was used to align both structures (RBD align), demonstrating changes in Nb6 and mNb6 position and the adjacent RBD. **E**, Comparison of mNb6 complementarity determining regions in either the cryo-EM structure of the Spike*-mNb6 complex or an X-ray crystal structure of mNb6 alone. **F**, CDR1 of Nb6 and mNb6 binding to the RBD. As compared to I27 in Nb6, Y27 of mNb6 hydrogen bonds to Y453 and optimizes pi-pi and pi-cation interactions with the RBD. **G**, CDR3 of Nb6 and mNb6 binding to the RBD demonstrating a large conformational rearrangement of the entire loop in mNb6. **H**, Comparison of closed Spike* bound to mNb6 and Nb6. Rotational axis for RBD movement is highlighted. **I**, Inhibition of pseudotyped lentivirus infection of ACE2 expressing HEK293T cells by mNb6 and mNb6-tri. n = 3 biological replicates **J**, mNb6 and mNb6-tri inhibit SARS-CoV-2 infection of VeroE6 cells in a plaque assay. Representative biological replicate with n = 4 technical replicates per concentration. n = 3 biological replicates for all samples. All error bars represent s.e.m.

**Figure 5.**
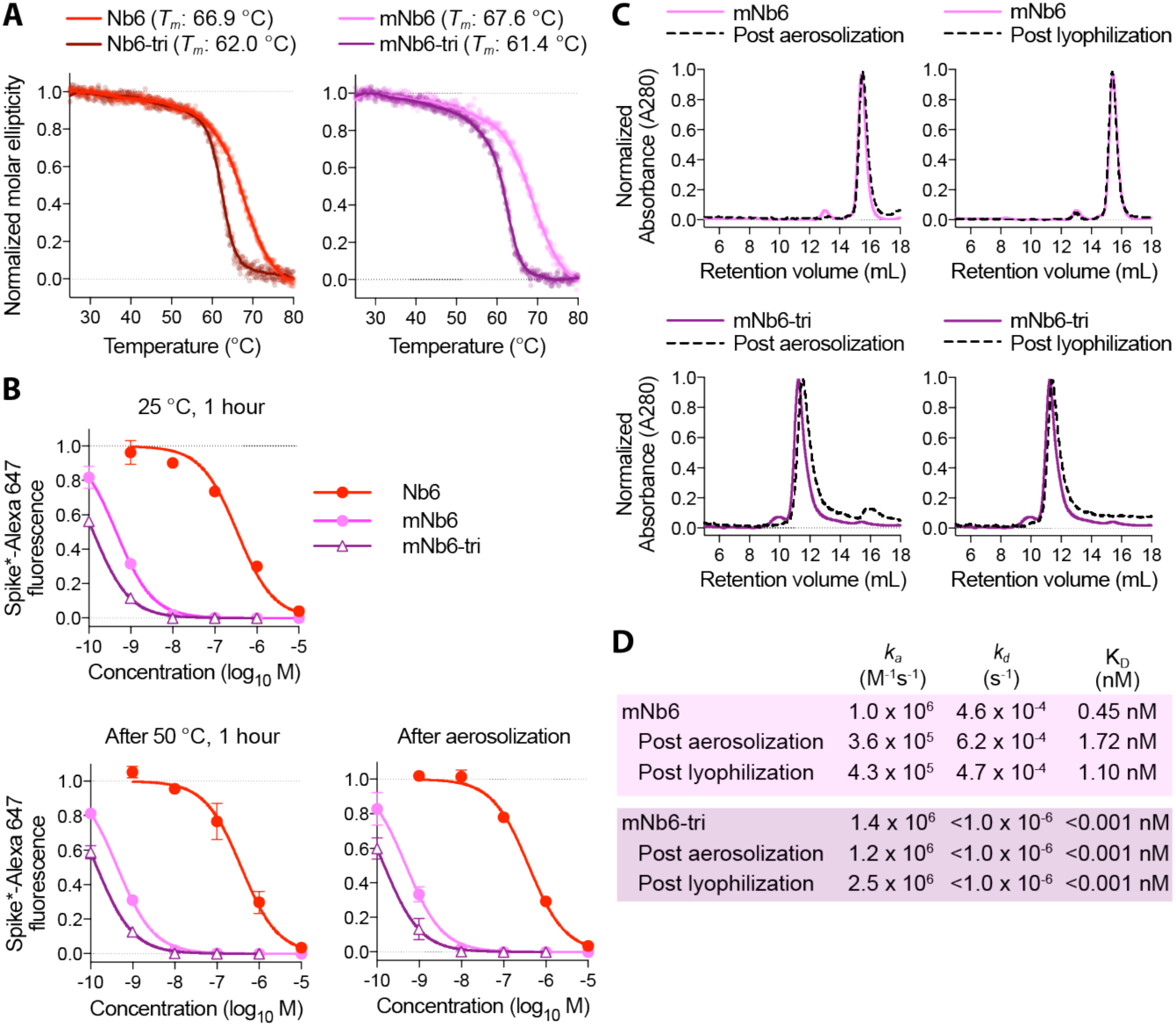
Nb6 and its derivates are robust proteins. **A**, Thermal denaturation of nanobodies assessed by circular dichroism measurement of molar ellipticity at 204 nm. Apparent melting temperatures (*T*_*m*_) for each nanobody are indicated. **B**, Nanobody inhibition of 1 nM Spike*-Alexa 647 binding to ACE2 expressing HEK293T cells after incubation at either 25 °C or 50 °C for 1 hour or after aerosolization. **C**, Size exclusion chromatography of nanobodies after lyophilization or aerosolization. **D**, Summary table of SPR kinetics data and affinities for aerosolized or lyophilized mNb6 and mNb6-tri.

A 2.9 Å cryo-EM structure of mNb6 bound to Spike* shows that, like the parent nanobody Nb6, mNb6 binds to closed Spike (Fig. 4C, Supplementary Fig. 7). The higher resolution map allowed us to build a model with high confidence and determine the effects of the I27Y and P105Y substitutions. mNb6 induces a slight rearrangement of the down-state RBDs as compared to both previously determined structures of apo-Spike* and Spike* bound to Nb6, inducing a 9° rotation of the RBD away from the central three-fold symmetry axis (Fig. 4H) (*14, 15*). This deviation likely arises from a different interaction between CDR3 and Spike*, which nudges the RBDs into a new resting position. While the I27Y substitution optimizes local contacts between CDR1 in its original binding site on the RBD, the P105Y substitution leads to a marked rearrangement of CDR3 in mNb6 (Fig. 4F-G). This conformational change yields a different set of contacts between mNb6 CDR3 and the adjacent RBD (Fig. 4D). Remarkably, an X-ray crystal structure of mNb6 alone revealed dramatic conformational differences in CDR1 and CDR3 between free and Spike*-bound mNb6, suggestive of significant conformational heterogeneity for the unbound nanobodies and induced-fit rearrangements upon binding to Spike* (Fig. 4E).

The binding orientation of mNb6 is similar to that of Nb6, supporting the notion that our multivalent design would likewise enhance binding affinity. Unlike Nb6-tri, trivalent mNb6 (mNb6-tri) bound to Spike with no observable fast-phase dissociation and no measurable dissociation over ten minutes, yielding an upper bound for the dissociation rate constant *k*_*d*_ of 1.0×10^−6^ s^-1^ (t_1/2_ > 8 days) and a K_D_ of <1 pM (Fig. 4B). As above, more precise measurements of the dissociation rate are precluded by the surface chemistry used to immobilize Spike*.

mNb6-tri displays further gains in potency in both pseudovirus and live SARS-CoV-2 infection assays with IC_50_ values of 120 pM (5.0 ng/mL) and 54 pM (2.3 ng/mL), respectively (Fig. 4H-I, Table 1). Given the sub-picomolar affinity observed by SPR, it is likely that these viral neutralization potencies reflect the lower limit of the assays. mNb6-tri is therefore an exceptionally potent SARS-CoV-2 neutralizing antibody, among the most potent molecules disclosed to date.

### Nb6, Nb6-tri, mNb6, and mNb6-tri are robust proteins

One of the most attractive properties that distinguishes nanobodies from traditional monoclonal antibodies is their extreme stability (*21*). We therefore tested Nb6, Nb6-tri, mNb6, and mNb6-tri for stability regarding temperature, lyophilization, and aerosolization. Temperature denaturation experiments using circular dichroism measurements to assess protein unfolding revealed melting temperatures of 66.9, 62.0, 67.6, and 61.4 °C for Nb6, Nb6-tri, mNb6 and mNb6-tri, respectively (Fig 5A). Aerosolization and prolonged heating of Nb6, mNb6, and mNb6-tri for 1 hour at 50°C induced no loss of activity (Fig 5B). Moreover, mNb6 and mNb6-tri were stable to lyophilization and to aerosolization using a mesh nebulizer, showing no aggregation by size exclusion chromatography and preserved high affinity binding to Spike* (Fig. 5C-D).

## DISCUSSION

There is a pressing need for prophylactics and therapeutics against SARS-CoV-2 infection. Most recent strategies to prevent SARS-CoV-2 entry into the host cell aim at blocking the ACE2-RBD interaction. High-affinity monoclonal antibodies, many identified from convalescent patients, are leading the way as potential therapeutics (*22-29*). While highly effective *in vitro*, these agents are expensive to produce by mammalian cell expression and need to be intravenously administered by healthcare professionals (*30*). Moreover, large doses are likely to be required for prophylactic viral neutralization, as only a small fraction of systemically circulating antibodies cross the epithelial cell layers that line the airways (*31*). By contrast, single domain antibodies (nanobodies) provide significant advantages in terms of production and deliverability. They can be inexpensively produced at scale in bacteria *(E. coli)* or yeast *(P. pastoris)*. Furthermore, their inherent stability enables aerosolized delivery directly to the nasal and lung epithelia by self-administered inhalation (*32*).

Monomeric mNb6 is among the most potent single domain antibodies neutralizing SARS-CoV-2 discovered to date. Multimerization of single domain antibodies has been shown to improve target affinity by avidity (*32, 33*). In the case of Nb6 and mNb6, however, our design strategy enabled a multimeric construct that simultaneously engages all three RBDs, yielding profound gains in potency. Furthermore, because RBDs must be in the up-state to engage with ACE2, conformational control of RBD accessibility can serve as an added neutralization mechanism. Indeed, our Nb6-tri and mNb6-tri molecules were designed with this functionality in mind. Thus, when mNb6-tri engages with Spike, it prevents ACE2 binding by both directly occluding the binding site and by locking the RBDs into an inactive conformation. Although a multitude of other monoclonal and single-domain antibodies against SARS-CoV-2 Spike have been discovered to date, there are few if any molecules as potent and stable as mNb6-tri (*33-43*). Resistance to aerosolization, in particular, offers unprecedented opportunity for patient-friendly nasal and pulmonary administration.

Our discovery of Class II neutralizing nanobodies demonstrates the presence of previously unexplored mechanisms of blocking Spike binding to ACE2. For one Class II nanobody, Nb3, we identified a likely binding site in the Spike S_1_ domain external to the RBDs. Previously discovered neutralizing antibodies from convalescent patients bind an epitope in a similar region of Spike (*24, 26, 27*). Binding of Nb3 to this epitope may allosterically stabilize RBDs in the down-state, thereby decreasing ACE2 binding. Pairing of Class I and Class II nanobodies in a prophylactic or therapeutic cocktail could thus be a highly advantageous strategy for both potent neutralization and prevention of escape variants. The combined stability, potency, and diverse epitope engagement of our anti-Spike nanobodies therefore provide a unique potential prophylactic and therapeutic strategy to limit the continued toll of the COVID-19 pandemic.

## ACKNOWLEDGEMENTS

We thank the entire Walter and Manglik labs for facilitating the development and rapid execution of this large-scale collaborative effort. We thank Sebastian Bernales and Tony De Fougerolles for advice and helpful discussion, and Jonathan Weissman for input into the project and reagent and machine use. We thank Jim Wells for providing the ACE2 ECD-Fc construct, Jason McLellan for providing Spike, RBD, and ACE2 constructs, and Florian Krammer for providing an RBD construct. We thank Jesse Bloom for providing the ACE2 expressing HEK293T cells as well as the plasmids for pseudovirus work. We thank George Meigs and other Beamline staff at ALS, 8.3.1 for their help in data collection. We thank Randy A. Albrecht for oversight of the conventional BSL3 biocontainment facility at the Icahn School of Medicine at Mount Sinai.

## Author Contributions

M.Schoof purified Spike*, RBD, and ACE2 proteins, performed yeast display selections to identify and affinity mature nanobodies, expressed and purified nanobodies, tested activity in cell-based assays, cloned, expressed, and purified multivalent nanobody constructs, and coordinated live virus experiments. B.Faust purified and characterized Spike* protein and candidate nanobodies, developed, performed and analyzed SPR experiments for Spike* and RBD-nanobody affinity determination, developed, performed and analyzed SPR binning, experiments, determined optimal freezing conditions for cryo-EM experiments, processed, refined and generated figures for Nb6, Nb11, and mNb6 EM datasets. R.A. Saunders expressed and purified ACE2 and nanobodies, developed and performed cell-based assays for inhibition of Spike* binding and pseudovirus assays for determining nanobody efficacy. S.Sangwan expressed and purified Spike*, RBD, ACE2-Fc, and nanobodies, processed cryo-EM data, optimized RBD-nanobody complexes for crystallography, grew crystals of mNb6, collected diffraction data, and refined the X-ray crystal structure of mNb6. V.Rezelj tested efficacy of nanobody constructs in live SARS-CoV-2 infection assays under the guidance of M.Vignuzzi. N.Hoppe purified nanobodies, developed, performed and analyzed SPR binning experiments, developed performed and analyzed variable Nb6-bi and Nb6-tri association experiments, and performed thermal melting stability assays for nanobody constructs. M.Boone developed approaches to express and purify nanobodies from *Pichia pastoris* and developed, performed, and analyzed approaches to quantify nanobody efficacy in live virus assays. C.B.Billesbølle expressed and purified Spike*, generated affinity maturation library for Nb6, performed yeast display selections to identify mNb6, and built the synthetic yeast nanobody library with J.Liang. I.Deshpande expressed and purified nanobody constructs. B.S.Zha performed live SARS-CoV-2 virus assays to test nanobody efficacy with guidance from O.Rosenberg. C.R.Simoneau and K.Leon performed live SARS-CoV-2 virus assays to test nanobody efficacy with guidance from M.Ott. K.M.White performed live SARS-CoV-2 virus assays to test nanobody efficacy with guidance from A.Garcia-Sastre. A.W.Barile-Hill performed SPR experiments. A.A.Anand, N.Dobzinski, B.Barsi-Rhyne, and Y.Liu. assisted in cloning, expression, and purification of nanobody and pseudovirus constructs. V.Belyy performed single-molecule nanobody-Spike* interaction studies. S.Nock prepared media and coordinated lab usage during UCSF’s partial shutdown. M.Zimanyi and S.Gupta performed radiolytic footprinting experiments with guidance from C.Y.Ralston and analyzed mass spectrometry data generated by D.L.Swaney. Several members of the QCRG Structural Biology Consortium played an exceptionally important role for this project. C.Azumaya and C.Puchades determined optimal freezing conditions for cryo-EM experiments, optimized data collection approaches, and collected cryo-EM datasets. A.F.Brilot, A.Rizo, A.M.Smith, F.Moss, D.Bulkley, T.Popsiech collected cryo-EM data on Spike*-nanobody complexes. S.Dickinson, H.C.Nguyen, C.M.Chio, U.S.Chio, M.Gupta, M.Jin, F.Li, Y.Liu, G.E.Merz, K.Zhang, M.Sun analyzed cryo-EM data from 15 Spike*-nanobody complex datasets. H.T.Kratochvil set up crystallization trials of various RBD-nanobody complexes, and crystallized, collected diffraction data for, and refined the mNb6 structure. M.C.Thompson collected, processed, and refined the mNb6 structure. R.Trenker, D.Diwanji, K.Schaefer expressed and purified Spike*, and S.Pourmal purified RBD. A.Manglik expressed and purified Spike*, labeled Spike* for biochemical studies, designed selection strategies for nanobody discovery, cloned nanobodies for expression, designed affinity maturation libraries and performed selections, analyzed SPR data, and performed nanobody stability studies. The overall project was supervised by P.Walter and A.Manglik.

## Data and Materials Availability

All data generated or analyzed during this study are included in this published article and its Supplementary Information. Crystallographic coordinates and structure factors for mNb6 have been deposited in the Protein Data Bank under accession code XXXX. Coordinates for Spike*:Nb6 and Spike*:mNb6 complexes have been deposited in the Protein Data Bank under accession codes XXXX and XXXX, respectively. Maps for Spike*:Nb6, Spike*:Nb11, and Spike*:mN6 have been deposited in the Electron Microscopy Data Bank under accession codes XXXXX (Spike*-Nb6 Open), XXXXX (Spike*-Nb6 Closed), XXXXX (Spike*-Nb11 Open), XXXXX (Spike*-Nb11 Closed), XXXXX (Spike*-mNb6 Closed). Plasmids for nanobody constructs used in this study are available under a Material Transfer Agreement with the University of California, San Francisco.

