## Supplementary_Information for "An ultra-potent synthetic nanobody neutralizes SARS-CoV-2 by locking Spike into an inactive conformation"

### SUPPLEMENTARY INFORMATION for Schoof *et al.*

#### MATERIALS AND METHODS

##### Expression and purification of SARS-CoV-2 Spike, RBD, and ACE2.

We used a previously described construct to express and purify the pre-fusion SARS-CoV-2 Spike ectodomain (Spike*) (*15*). ExpiCHO or Expi293T cells (ThermoFisher) were transfected with the Spike* construct per the manufacturer's instructions for the MaxTiter protocol and harvested between 3-9 days after transfection. Clarified cell culture supernatant was loaded onto Ni-Excel beads (Cytiva) followed by extensive washes in 20 mM HEPES pH 8.0, 200 mM sodium chloride, and 10 mM imidazole and elution in the same buffer supplemented with 500 mM imidazole. Spike* was concentrated using a 100 kDa MWCO spin concentrator (Millipore) and further purified by size exclusion chromatography over a Superose 6 Increase 10/300 column (GE Healthcare) in 20 mM HEPES pH 8.0 and 200 mM sodium chloride. All purification steps were performed at room temperature. The resulting fractions for trimeric Spike* were pooled and either used directly for cryo-EM studies or concentrated and flash frozen in liquid nitrogen with 15% glycerol for other biochemical studies.

We used a previously described construct to express and purify the SARS-CoV-2 Receptor binding domain (RBD) (*44*). Expi293T cells (ThermoFisher) were transfected with the RBD construct per the manufacturer's instructions and harvested between 3-6 days after transfection. Clarified cell culture supernatant was loaded onto Ni-Excel beads (Cytiva) or a His-Trap Excel column (GE Healthcare) followed by washes in 20 mM HEPES pH 8.0, 200 mM sodium chloride, and 10 mM imidazole and elution using the same buffer supplemented with 500 mM imidazole. RBD was concentrated using a 30 kDa MWCO spin concentrator (Millipore) and further purified by size exclusion chromatography over a Superdex 200 Increase 10/300 GL column (GE Healthcare) in 20 mM HEPES pH 8.0 and 200 mM sodium chloride. The resulting fractions were pooled, concentrated, and flash frozen in liquid nitrogen with 10% glycerol.

For biochemical and yeast display experiments, Spike* and RBD were labeled with freshly prepared stocks of Alexa 647-NHS, Alexa 488-NHS, or Biotin-NHS (ThermoFisher) with a 5-fold stoichiometry for 1 hour at room temperature followed by quenching of NHS with 10 mM Tris pH 8.0 for 60 minutes. Labeled proteins were further purified by size exclusion chromatography, concentrated using a spin concentrator (Millipore), and flash frozen in liquid nitrogen with 10-15% glycerol.

We used an ACE2-ECD (18-614) Fc fusion expression plasmid to express and purify Fc tagged ACE2-ECD (*45*). Expi293T cells (ThermoFisher) were transfected with the ACE2-Fc construct per the manufacturer's instructions and harvested between 5-7 days after transfection. Clarified cell culture supernatant was loaded onto a MabSelect Pure 1 mL Column (GE Healthcare). Column was washed with Buffer A (20 mM HEPES pH 7.5, 150 mM NaCl) and protein was eluted with Buffer B (100 mM Sodium Citrate pH 3.0, 150 mM NaCl) into a deep well block containing 1 M HEPES pH 7.5 to neutralize the acidic elution. ACE2-Fc was concentrated using a 30 kDa MWCO spin concentrator (Millipore) and further purified by size exclusion chromatography over a Superdex 200 Increase 10/300 GL column (GE Healthcare) in SEC Buffer (20 mM HEPES pH 7.5, 150 mM NaCl, 5% v/v Glycerol). The resulting fractions were pooled, concentrated, and flash frozen in liquid nitrogen. To obtain monomeric ACE2, 1:50 (w/w) His-tagged TEV protease was added to ACE2-Fc and incubated at 4 °C overnight. This mixture was then purified by size exclusion chromatography in SEC Buffer. Monomeric ACE2 fractions were pooled and washed with His-resin (1 mL of 50% slurry) to remove excess TEV. The resulting supernatant was pooled, concentrated, and flash frozen in liquid nitrogen.

**Identification of anti SARS-CoV2 Spike nanobodies**

To identify nanobodies against the SARS-CoV-2 Spike ECD, we used a yeast surface displayed library of synthetic nanobody sequences that recapitulate amino acid position specific-variation in natural llama immunological repertoires. This library encodes a diversity of >$2x10^9$ variants, and uses a synthetic stalk sequence for nanobody display, as described previously in a modified vector encoding nourseothricin (NTC) resistance (*46*). For the first round of selection, $2x10^{10}$ yeast induced in YPG (Yeast Extract-Peptone-Galactose) supplemented with NTC were washed repeatedly in selection buffer (20 mM HEPES, pH 7.5, 150 mM sodium chloride, 0.1% (w/v) low biotin bovine serum albumin, BSA) and finally resuspended in 10 mL of selection buffer containing 200 nM biotinylated-Spike*. Yeast were incubated for 30 minutes at 25 °C, then washed repeatedly in cold selection buffer, and finally resuspended in 10 mL of cold selection buffer containing 200 µL of Miltenyi anti-Streptavidin microbeads. After 30 minutes of incubation at 4 °C, yeast were again washed with cold selection buffer. Spike* binding yeast were captured on a Miltenyi MACS LS column and recovered in YPD (Yeast Extract-Peptone-Dextrose) medium supplemented with NTC.

For round 2, $4x10^8$ induced yeast from Round 1 were incubated with 100 nM Spike* labeled with Alexa647 in 1 mL of selection buffer for 1 hr at 25 °C. After extensive washes with cold selection buffer, Spike* binding yeast were isolated by fluorescence activated cell sorting (FACS) on a Sony SH800 instrument. A similar approach was used for round 3, with substitution of 10 nM Spike* labeled with Alexa647. Post round 3 yeast were plated on YPD+NTC solid media and 768 individual colonies were induced with YPG+NTC media in 2 mL deep well plates. Each individual clone was tested for binding to 4 nM Spike*-Alexa488 by flow cytometry on a Beckman Coulter Cytoflex. To identify nanobodies that disrupt Spike-ACE2 interactions, Spike* binding was repeated in the presence of 0.5-1 µM ACE2-Fc. Out of 768 clones, we identified 21 that strongly bind Spike* and are competitive with ACE2 (Supplementary Table 3).

**Expression and purification of nanobodies**

Nanobody sequences were cloned into the pET26-b(+) expression vector using In-Fusion HD cloning (Takara Bio), transformed into BL21(DE3) *E. coli*, grown in Terrific Broth at 37 °C until OD 0.7-0.8, followed by gene induction using 1 mM IPTG for 18-22 hours at 25°C. *E. Coli* were harvested and resuspended in SET Buffer (200 mM Tris, pH 8.0, 500 mM sucrose, 0.5 mM EDTA, 1X cOmplete protease inhibitor (Roche)) for 30 minutes at 25 °C before a 45 minute osmotic shock with a two-fold volume addition of water. NaCl, MgCl2, and imidazole were added to the lysate to 150 mM, 2 mM, and 40 mM respectively before centrifugation at 17-20,000xg for 15 minutes to separate cell debris from the periplasmic fraction. For every liter of bacterial culture, the periplasmic fraction was then incubated with 4 mL of 50% HisPur Ni-NTA resin (Thermo Scientific) which had been equilibrated in Nickel Wash Buffer (20 mM HEPES, pH 7.5, 150 mM NaCl, 40 mM imidazole). This mixture was incubated for 1 hr with rotation at RT before centrifugation at 50xg to collect the resin. The resin was then washed with 5 volumes of Nickel Wash buffer 3 times, each time using centrifugation to remove excess wash buffer. Bound proteins were then eluted using three washes with Elution Buffer (20 mM HEPES, pH 7.5, 150 mM NaCl, 500 mM imidazole). The eluted protein was concentrated using a 3.5 kDa MWCO centrifugal filter unit (Amicon) before injection onto a Superdex 200 Increase 10/300 GL column equilibrated with 20 mM HEPES, pH 7.5, 150 mM NaCl. Nanobody constructs were concentrated again using a 3.5k MWCO centrifugal filter unit, and flash frozen in liquid nitrogen.

**Affinity determination by surface plasmon resonance**

Nanobody (Nb) affinity determination experiments were performed on Biacore T200 and 8K instruments (Cytiva Life Sciences) by capturing the StreptagII-tagged Spike* at 10 µg/mL on a

StreptactinXT-immobilized (Iba Life Sciences) CM5 Series S sensor chip (Cytiva Life Sciences) to achieve maximum response (Rmax) of approximately 30 response units (RUs) upon nanobody binding. 2-fold serial dilutions of purified nanobody from 1 μM to 31.25 nM (for monovalent constructs) or from 50 nM to 1.56 nM (for affinity matured and multimeric constructs) were flowed over the captured Spike* surface at 30 μL/minute for 60 seconds followed by 600 seconds of dissociation flow. Following each cycle, the chip surface was regenerated with 3 M guanidine hydrochloride.

Separately, biotinylated SARS-CoV-2 RBD at 8 μg/mL was loaded onto a preconditioned Series S Sensor Chip CAP chip (Cytiva Life Sciences) to achieve an Rmax of approximately 60 RUs upon nanobody binding. 2-fold serial dilutions in the same running buffer and sample series (parent or affinity matured clone) as the Spike* runs were flowed over the RBD surface at 30 μL/minute for 60 seconds followed by 600 seconds of dissociation flow. Chip surface regeneration was performed with a guanidine hydrochloride/sodium hydroxide solution.

The resulting sensorgrams for all monovalent clones were fit to a 1:1 Langmuir binding model using the Biacore Insight Evaluation Software (Cytiva Life Sciences) or the association/dissociation model in GraphPad Prism 8.0. For determination of kinetic parameters for Nb6-bi and Nb6-tri binding, the dissociation phase was fit to a biexponential decay constrained to two dissociation rate constants shared between each concentration. The association phase was fit separately using an association kinetics model simultaneously fitting the association rate constant for each concentration.

For nanobody competition experiments, Spike* was loaded onto a StreptactinXT-immobilized CM5 sensor chip as previously described. As in the kinetics experiments, the primary nanobody was flowed over the captured Spike* surface for 60 seconds at 30 μL/minute to achieve saturation. Immediately following this, a second injection of a mixture of primary and variable nanobody at the same concentration as in the primary injection was performed.

**ACE2 cellular surface binding competition assays**

A dilution series of nanobody was generated in PBE (PBS + 0.5% (w/v) BSA + 2 mM EDTA and mixed with Spike*-Alexa647 or RBD-Alexa647. ACE2 expressing HEK293T cells were dissociated with TrypLE Express (ThermoFisher) and resuspended in PBE (*20*). The cells were mixed with the Spike*-nanobody solution and incubated for 45 minutes, washed in PBE, and then resuspended in PBE. Cell surface Alexa647 fluorescence intensity was assessed on an Attune Flow Cytometer (ThermoFisher).

**Affinity maturation of Nb6**

A site saturation mutagenesis library of Nb6 was generated by assembly PCR of overlapping oligonucleotides encoding the Nb6 sequence. Individual oligos for each position in CDR1, CDR2, and CDR3 were designed with the degenerate “NNK” codon. The assembled gene product was amplified with oligonucleotides with overlapping ends to enable homologous recombination with the yeast surface display vector as previously described and purified with standard silica-based chromatography (*46*). The resulting insert DNA was transformed into *Saccharomyces cerevisiae* strain BJ5465 along with the yeast display vector pYDS2.0 to generate a library of $2x10^8$ transformants. After induction in YPD+NTC medium at 20 °C for 2 days, $2x10^9$ yeast were washed in selection buffer (20 mM HEPES, pH 8.0, 150 mM sodium chloride, 0.1% (w/v) low biotin BSA) and incubated with 1 nM biotin-Spike* for 1 hour at 25 °C. Yeast were subsequently washed in selection buffer, resuspended in 1 mL selection buffer, and incubated with 10 µL streptavidin microbeads (Miltenyi) for 15 min. at 4 °C. Yeast were washed again with cold selection buffer and Spike*-binding yeast were isolated by magnetic separation using an LS column (Miltenyi). Recovered yeast were grown in YPD+NTC at 37 °C and induced in YPG+NTC at 20 °C. A second round of selection was performed as above, substituting 100 pM RBD-Alexa647 as the antigen. Yeast displaying high affinity clones were selected by magnetic separation using Anti-Cy5 microbeads (Miltenyi) and an LS column. Analysis of the library after the second round of selection revealed a population of clones with clear binding of 10 pM RBD-Alexa647. Therefore, 96 individual clones were screened for binding to 10 pM RBD-Alexa647 by flow cytometry. Sequence analysis of eight clones that showed robust binding to 10 pM RBD-Alexa647 revealed two consensus mutations, I27Y and P105Y, which were used to generate the affinity matured clone mNb6.

**Structures of Spike-nanobody complexes by cryo-EM**

*Sample preparation and microscopy*

To prepare Spike*-nanobody complexes, each nanobody was incubated on ice at a 3-fold molar excess to Spike* at 2.5 µM for 10 minutes. 3 µL of Spike*-nanobody complex was added to a 300 mesh 1.2/1.3R Au Quantifoil grid previously glow discharged at 15 mA for 30 seconds. Blotting was performed with a blot force of 0 for 4 seconds at 4°C and 100% humidity in a FEI Vitrobot Mark IV (ThermoFisher) prior to plunge freezing into liquid ethane.

For each complex, 120-frame super-resolution movies were collected with a 3x3 image shift collection strategy at a nominal magnification of 105,000x (physical pixel size: 0.834 Å/pix) on a Titan Krios (ThermoFisher) equipped with a K3 camera and a Bioquantum energy filter (Gatan) set to a slit width of 20 eV. Collection dose rate was 8 $e^-$/pixel/second for a total dose of 66 $e^-$/$Å^2$. Each collection was performed with semi-automated scripts in SerialEM (*47*).

*Image Processing*

For all datasets, dose fractionated super-resolution movies were motion corrected with MotionCor2 (*48*). Contrast transfer function determination was performed with cryoSPARC patch CTF (*49*). Particles were picked with a 20 Å low-pass filtered apo Spike 2D templates generated from a prior data collection.

Nb6-Spike* and mNb6-Spike* particles were extracted with a 384 pixel box, binned to 96 pixels and subject to single rounds of 2D and 3D classification prior to unbinning for homogenous refinement in cryoSPARC (*49*). Refined particles were then imported into Relion3.1 for 3D classification without alignment using the input refinement map low pass filtered to 40 Å (*50*). Particles in classes representing the closed conformation of Spike were imported into cisTEM and subject to autorefinement followed by local refinement within a RBD::nanobody masked region (*51*). Following local refinement, a new refinement package symmetrized to the C3 axis was created for a final round of local refinement without masking. Final particle counts for each map are as follows: Nb6-Open: 40,125, Nb6-Closed: 58,493, mNb6: 53,690.

Nb11-Spike* particles were extracted with a 512 pixel box, binned to 128 pixels for multiple rounds of 3D classification as described in Figure S4. Following homogenous refinement, particles were exported to Relion3.1. Particle density roughly corresponding to RBD-nanobody complexes was retained post-particle subtraction. 3D classification without alignment was performed on the particle subtracted stacks. Particles in classes with robust RBD-nanobody density were selected, unsubtracted and refined in Relion followed by post-processing. 21,570 particles contributed to the final maps. Final particle counts for each map are as follows: Nb11-Open: 21,570, Nb11-Closed: 27,611. For all maps, final local resolution estimation and GSFSC determination was carried out in cryoSPARC.

*Structure modeling*

Models of Nb6-Spike* and mNb6-Spike* were built using a previously determined structure of closed Spike* (PDB: 6VXX) (*14*). A composite model incorporating resolved regions of the RBD was made using a previously determined X-ray crystal structure of the SARS-CoV-2 RBD (PDB: 6M0J) (*52*). For Nb6, the beta2-adrenergic receptor nanobody Nb80 (PDB: 3P0G) was used as a template to first fit the nanobody into the cryo-EM density map for the Nb6-Spike* complex (*53*). Complementarity determining loops were then truncated and rebuilt using RosettaES (*54*). The final structure was inspected and manually adjusted in COOT and ISOLDE, followed by real space refinement in PHENIX (*55*-*57*). The higher resolution structure of mNb6 enabled manual building of nanobody CDR loops *de novo*, and therefore the Rosetta-based approach was not used for modeling. Final models were analyzed in PHENIX, with statistics reported in Supplementary Table 1.

For models of Nb11-Spike* complexes presented here, the closest nanobody by sequence in the PDB (beta2-adrenergic receptor Nb60, PDB ID: 5JQH) was fit by rigid-body refinement in COOT into the cryo-EM density map using only the framework regions (*58*). While the lower resolution of these maps precluded confident assignment of loop conformations, the overall orientation of Nb11 relative to Spike* was well constrained, enabling accurate modeling of distances between the N- and C- termini of two Nb11 molecules bound to Spike*.

**Radiolytic hydroxyl radical footprinting and mass-spectrometry of Spike* and Nb3-Spike***

Spike* and Nb3 samples were buffer exchanged into 10 mM phosphate buffer (pH 7.4) by extensive dialysis at 25 °C. A 1.5-fold molar excess of Nb3 was added to 5 µM Spike* and the complex was incubated for >24 hr at 25 °C. For radiolytic footprinting, protein concentrations and beam parameters were optimized using an Alexa-488 fluorophore assay (*59*). Apo Spike* and Spike*-Nb3 complex at concentrations of 1-3 µM were exposed to a synchrotron X-ray white beam at 6 timepoints between 0-50 ms at beamline 3.2.1 at the Advanced Light Source in Berkeley, CA and were quenched with 10 mM methionine amide immediately post-exposure. Glycans were removed by treatment with 5% SDS, 5 mM DTT at 95 °C for five minutes and subsequent PNGase (Promega) digestion at 37°C for 2 hours. Samples were buffer exchanged into ammonium bicarbonate (ABC) buffer (pH 8.0) using ZebaSpin columns (Thermo Fisher). Alkylation of cysteines was achieved by treatment with 8 M urea and 5 mM DTT at 37°C for 30 minutes followed by an incubation with 15 mM iodoacetamide at 25 °C in the dark for 30 minutes. All samples were further buffer exchanged to ABC pH 8.0 using ZebaSpin columns and digested with either Trypsin/Lys-C or Glu-C (Promega) at an enzyme:protein ratio of 1:20 (w/w) at 37 °C for 8 hours.

Samples were lyophilized and resuspended in 1% formic acid at 200 fmol/µL concentration. For each MS analysis, 1 µL of sample was injected onto a 5 mm Thermo Trap C18 cartridge, and then separated over a 15 cm column packed with 1.9 µm Reprosil C18 particles (Dr. Maisch HPLC GmbH) by a nanoElute HPLC (Bruker). Separation was performed at 50 °C and a flow rate of 400 µL/min by the following gradient in 0.1% formic acid: 2% to 17% acetonitrile from 0 to 20 min, followed by 17% to 28% acetonitrile from 20 to 40 min. The eluent was electrospray ionized into a Bruker timsTOF Pro mass spectrometer and data was collected using data-dependent PASEF acquisition. Database searching and extraction of MS1 peptide abundances was performed using the FragPipe platform with either trypsin or GluC enzyme specificity, and all peptide and protein identifications were filtered to a 1% false-discovery rate (*60*). Searches were performed against a concatenated protein database of the Spike protein, common contaminant proteins, and the *Saccharomyces cerevisiae* proteome (downloaded July 23, 2020). Note, the *Saccharomyces cerevisiae* proteome was included to generate a sufficient population of true negative identifications for robust false discovery rate estimation of peptide and protein identifications. Lastly, the area under the curve MS1 intensities reported from FragPipe were summarized for each peptide species using MSstats (*61*).

The peak areas of extracted ion chromatograms and associated side-chain modifications were used to quantify modification at each timepoint. Increasing beamline exposure time decreases the fraction of unmodified peptide and can be represented as a site-specific dose-response plot (Supplementary Fig. 5B). The rate of hydroxyl radical reactivity ($k_{fp}$) is dependent on both the intrinsic reactivity of each residue and its solvent accessibility and was calculated by fitting the dose-response to a pseudo-first order reaction scheme in Graphpad Prism Version 8. The ratio of $k_{fp}$ between apo Spike* and the Spike-Nb3 complex at specific residues gave information on solvent accessibility changes between the two samples. These changes were mapped onto the SARS-CoV-2 Spike (PDB 6XR8) (*11*). In some cases, heavily modified residues show a flattening of dose-response at long exposures which we interpret as radical induced damage. These over-exposed timepoints were excluded from the calculation of $k_{fp}$.

**mNb6 crystallography and structure determination**

Purified mNb6 was concentrated to 18.7 mg/mL and filtered using 0.1 µm hydrophilic PVDF filters (Millipore). mNb6 crystal screens were set up in 96 well plates in hanging drop format at 2:1 protein:reservoir in Index and AmSO4 screens (Hampton Research, Aliso Viejo, CA). Crystals in over 60 different screening conditions with various morphologies appeared overnight at ambient temperature and were obtained directly from the screens without further optimization. The crystals were cryoprotected by quick dipping in a solution containing 80% reservoir and 20% PEG400 or 20% Glycerol, then mounted in CrystalCap HT Cryoloops (Hampton Research, Aliso Viejo, CA) and flash cooled in a cryogenic nitrogen stream (100 K). All data were collected at the Advanced Light Source (Berkeley, CA) beam line 8.3.1. A single crystal of mNb6 that grew in 0.1 M Tris.HCl pH 8.5, 1.0 M Ammonium sulfate diffracted to 2.05 Å. Integration, and scaling were performed with Xia2, using XDS for indexing and integration and XSCALE for scaling and merging (*62*). The structure was solved molecular replacement using PHASER using the structure of nanobody, Nb.b201 (PDB 5VNV) as search model (*46, 63*). Model building was performed with COOT and refined with PHENIX and BUSTER(*55, 57, 64*).

**Pseudovirus assays for nanobody neutralization**

ZsGreen SARS-CoV-2-pseudotyped lentivirus was generated according to a published protocol (*20*). The day before transduction, 50,000 ACE2 expressing HEK293T cells were plated in each well of a 24-well plate. 10-fold serial dilutions of nanobody were generated in complete medium (DMEM + 10% FBS + PSG) and pseudotyped virus was added to a final volume of 200 µL. Media was replaced with nanobody/pseudotyped virus mixture for four hours, then removed. Cells were washed with complete medium and then incubated in complete medium at 37 °C. Three days post-transduction, cells were trypsinized and the proportion of ZsGreen+ cells was measured on an Attune flow cytometer (ThermoFisher).

**Authentic SARS-CoV-2 neutralization assay**

SARS-CoV-2, isolate France/IDF0372/2020, was supplied by the National Reference Centre for Respiratory Viruses hosted by Institut Pasteur (Paris, France) and headed by Pr. Sylvie van der Werf. Viral stocks were prepared by propagation in Vero E6 cells in Dulbecco's modified Eagle's medium (DMEM) supplemented with 2% (v/v) fetal bovine serum (FBS, Invitrogen). Viral titers were determined by plaque assay. All plaque assays involving live SARS-CoV-2 were performed at Institut Pasteur Paris (IPP) in compliance with IPP's guidelines following Biosafety Level 3 (BSL-3) containment procedures in approved laboratories. All experiments were performed in at least three biologically independent samples.

Neutralization of infectious SARS-CoV-2 was performed using a plaque reduction neutralization test in Vero E6 cells (CRL-1586, ATCC). Briefly, nanobodies (or ACE2-Fc) were eight-fold serially diluted in DMEM containing 2% (v/v) FBS and mixed with 50 plaque forming units (PFU) of SARS-CoV-2 for one hour at 37°C, 5% $CO_2$. The mixture was then used to inoculate Vero E6 cells seeded in 12-well plates, for one hour at 37 °C, 5% CO2. Following this virus adsorption time, a solid agarose overlay (DMEM, 10% (v/v) FBS and 0.8% agarose) was added. The cells were incubated for a further 3 days prior to fixation using 4% formalin and plaques visualized by the addition of crystal violet. The number of plaques in quadruplicate wells for each dilution was used to determine the half maximal inhibitory concentrations ($IC_{50}$) using 3-parameter logistic regression (GraphPad Prism version 8).

**Nanobody stability studies**

Nanobody thermostability by circular dichroism was assessed using a Jasco J710 CD spectrometer equipped with a Peltier temperature control. Individual nanobody constructs were diluted to 5 µM in phosphate buffered saline. Mollar ellipticity was measured at 204 nm (2 nm bandwidth) between 25 °C and 80 °C with a 1 °C/min heating rate. The resulting molar ellipticity values were normalized and plotted in GraphPad Prism 8.0 after applying a nearest neighbor smoothing function.

For nanobody competition experiments on ACE2 expressing HEK293T cells, nanobodies were incubated at either 25°C or 50°C for one hour. Alternatively, each nanobody was aerosolized with a portable mesh nebulizer producing 2-5 µm particles at a final concentration of 0.5 mg/mL. The resulting aerosol was collected by condensation into a 50 mL tube cooled on ice. Samples were then treated as indicated above to determine IC50 values for binding to Spike*-Alexa647.

Further experiments assessing mNb6 and mNb6-tri stability to aerosolization and lyophilization used a starting concentration of 0.5 mg/mL of each construct. Aerosolization was performed as described above. For lyophilization, nanobodies were first flash frozen in liquid nitrogen and the solution was dried to completion under vacuum. The resulting dried material was resuspended in 20 mM HEPES pH 7.5, 150 mM NaCl. Size exclusion chromatography of the unstressed, post-aerosolization, and post-lyophilization samples were performed an a Superdex 75 Increase 10/300 column in 20 mM HEPES pH 7.5, 150 mM NaCl. SPR experiments to assess binding to Spike* were performed as described above.

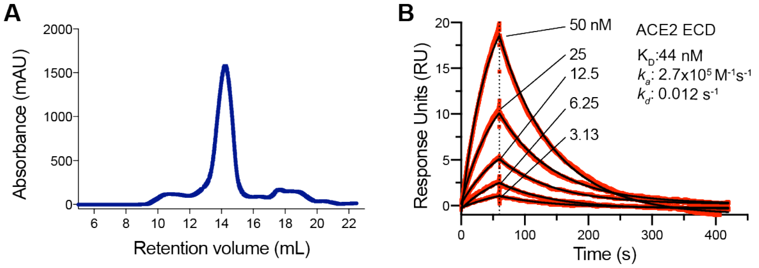

**Supplementary Fig. 1. Validation of purified Spike*. A,** Size exclusion chromatogram of purified Spike* from ExpiCHO cells. **B,** SPR of immobilized Spike* binding to monomeric ACE2 extracellular domain (ECD).

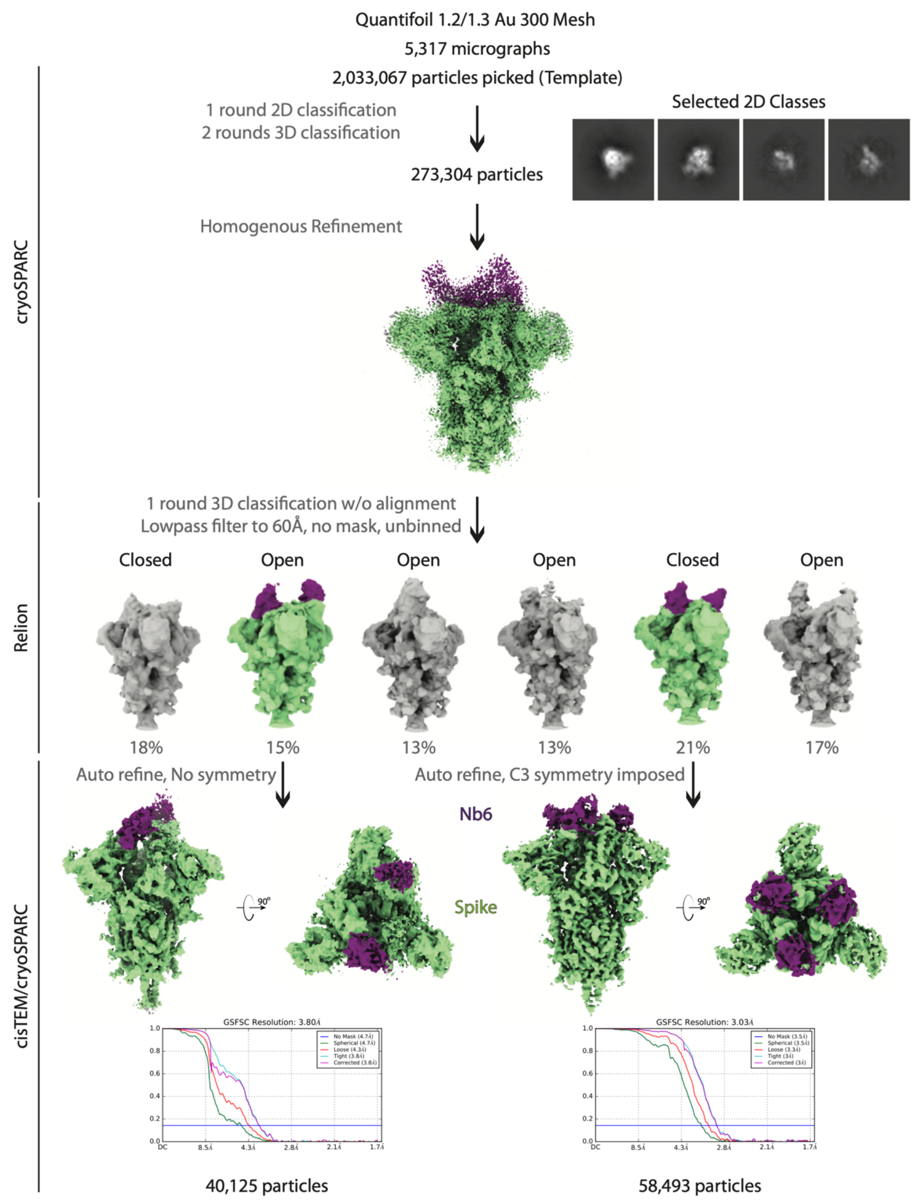

**Supplementary Fig. 2. Cryo-EM workflow for Nb6**

A flowchart representation of the classification workflow for Spike*-Nb6 complexes yielding open and closed Spike* conformations. From top to bottom, particles were template picked with a set of 20 Å low-pass filtered 2D backprojections of apo-Spike* in the closed conformation. Extracted particles in 2D classes suggestive of various Spike* views were subject to a round of heterogenous refinement in cryoSPARC with two naïve classes generated from a truncated *Ab initio* job, and a 20 Å low-pass filtered volume of apo-Spike* in the closed conformation. Particles in the Spike* 3D class were subject to 25 iterations of 3D classification into 6 classes without alignment in RELION, using the same input volume from cryoSPARC 3D classification, low pass filtered to 60 Å, T = 8. Particles in classes representing the open and closed Spike* conformations were imported into cisTEM for automatic refinement. Half maps from refinement were imported into cryoSPARC for local resolution estimation as shown in Supplementary Fig. 4.

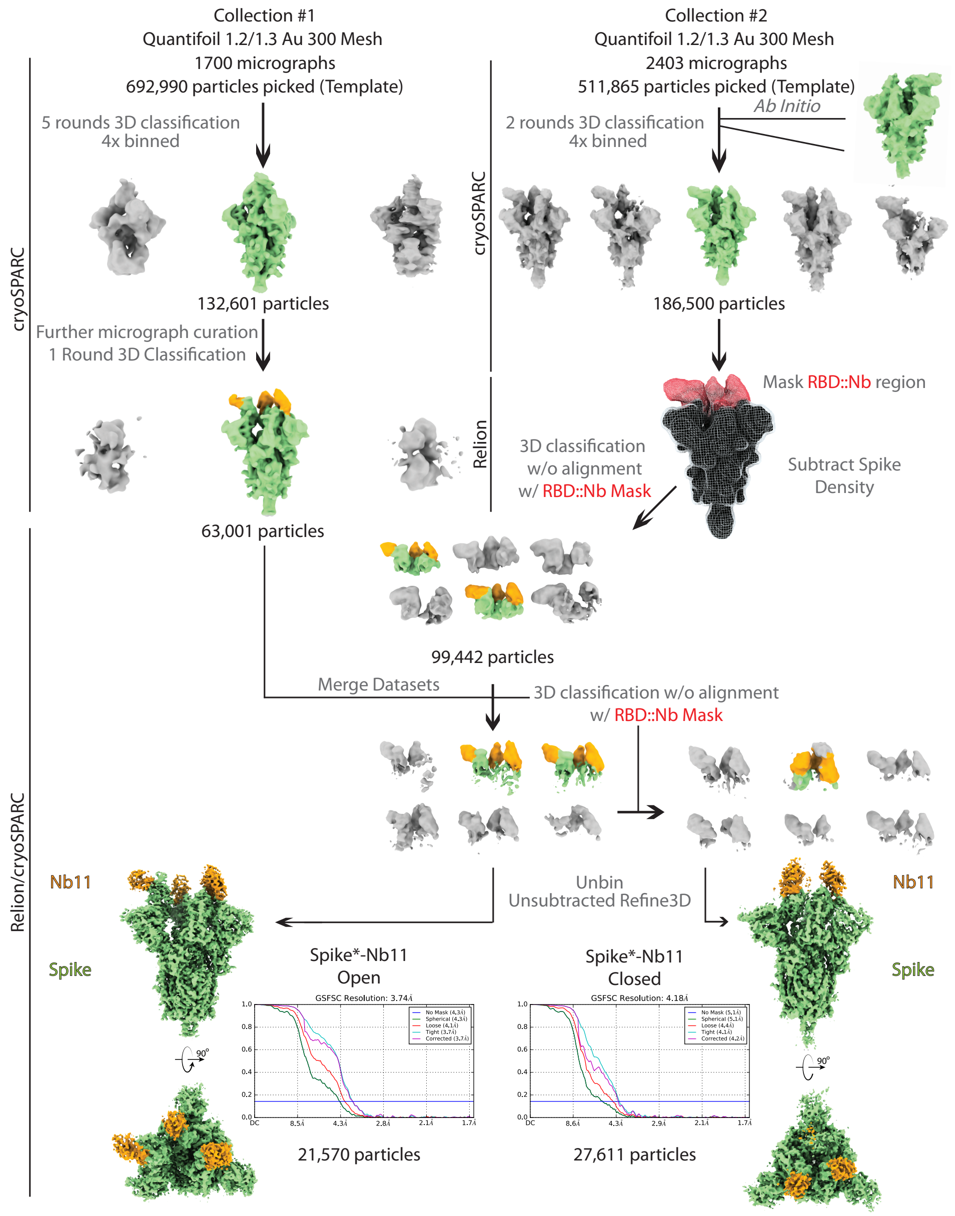

**Supplementary Fig. 3. Cryo-EM workflow for Nb11**

A flowchart representation of the classification workflow for Spike*-Nb11 complexes yielding open and closed Spike* conformations. From top to bottom, particles were template picked from two separate collections with a set of 20 Å low-pass filtered 2D backprojections of apo-Spike* in the closed conformation. Extracted particles were Fourier cropped to 128 pixels prior to extensive heterogenous refinement in cryoSPARC, using a 20 Å low-pass filtered volume of apo-Spike* in the closed conformation and additional naïve classes for removal of non-Spike* particles. After cryoSPARC micrograph curation and heterogenous refinement, Spike* density corresponding to all regions outside of the ACE2 RBD::Nanobody interface were subtracted. A mask around the ACE2 RBD::Nanobody interface was generated, and used for multiple rounds of 3D classification without alignment in RELION. Particles in classes representing open and closed Spike* conformations were selected, unsubtracted and unbinned prior to refinement in RELION. Half maps from refinement were imported into cryoSPARC for local resolution estimation as shown in Supplementary Fig. 4.

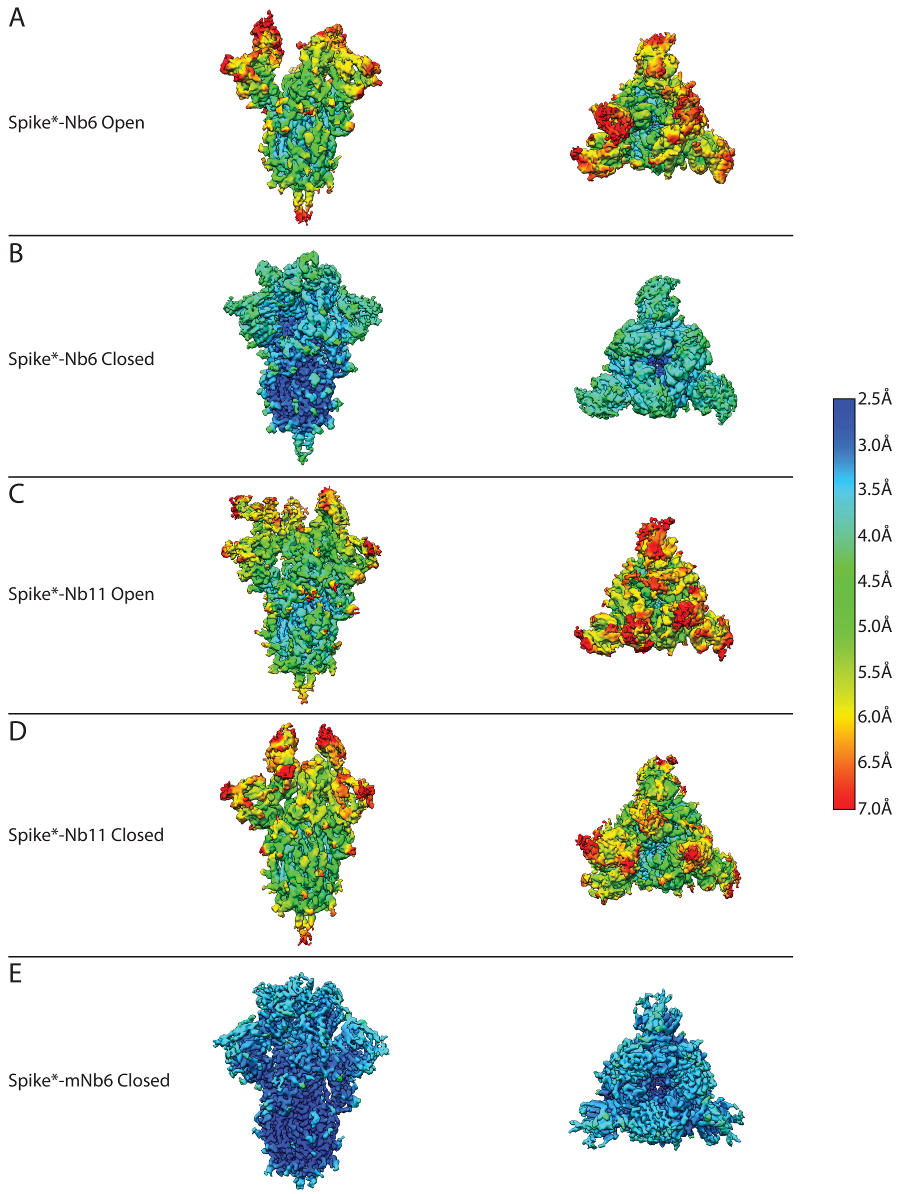

**Supplementary Fig. 4. Local resolution of cryo-EM maps**

Local resolution estimates of Spike* complexes with A-B) Nb6, C-D) Nb11, and E) mNb6 as generated in cryoSPARC. All maps (except mNb6) are shown with the same enclosed volume. All maps are colored on the same scale, as indicated.

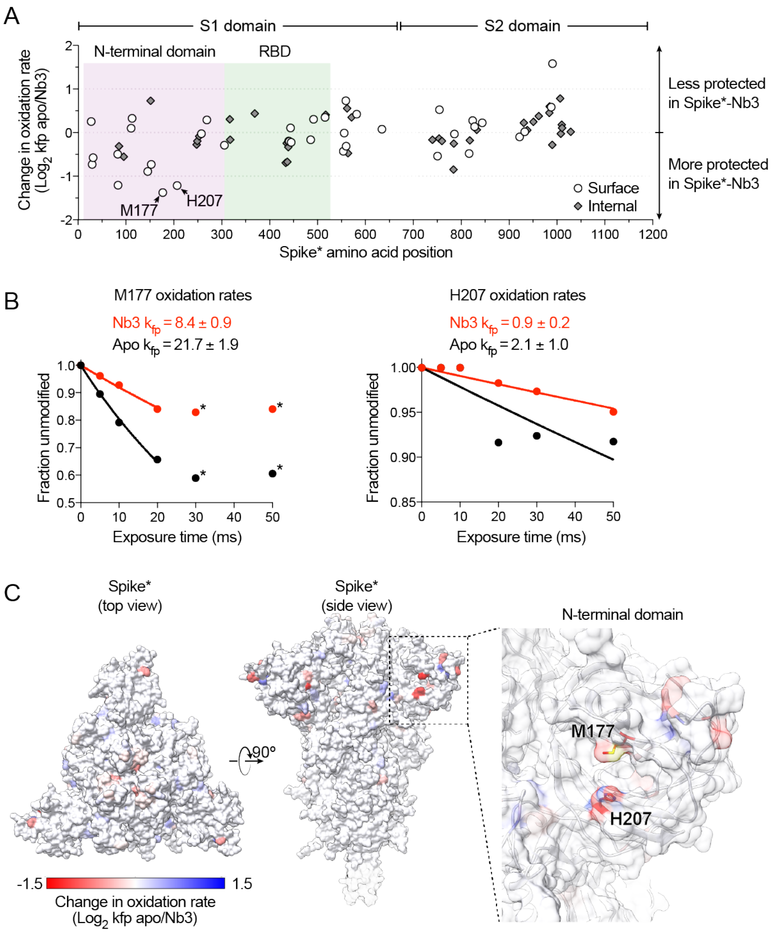

**Supplementary Fig. 5. Radiolytic hydroxyl radical footprinting of Spike*.**

**A,** Change in oxidation rate between Spike* and Nb3-Spike* complexes at all residues. A cluster of highly protected residues in the Spike*-Nb3 complex is observed in the N-terminal domain. **B,** Oxidation rate plots of the two (M177, H207) most heavily protected residues upon Nb3 binding to Spike*. Data points labeled with an asterisk are excluded from rate calculations as these values fall outside of the first order reaction, likely due to extensive oxidation-mediated damage. **C,** Change in oxidation rate mapped onto Spike in the all RBD down conformation.

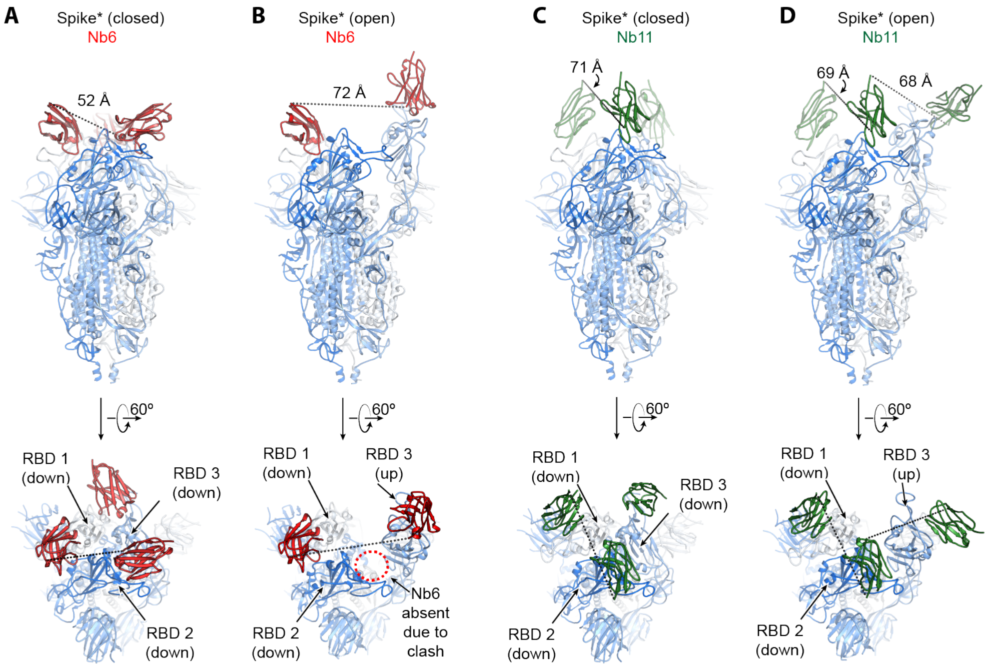

**Supplementary Fig. 6. Modeling of distances for multimeric nanobody design. A,** Model of Spike*:Nb6 complex in the closed state. The minimal distance between adjacent Nb6 N- and C-termini is 52 Å (dashed line). **B,** Model of Spike*:Nb6 complex in the open state with Nb6 docked into the cryo-EM density for up-state RBD. Minimal distance between N- and C-termini of both nanobodies is 72 Å**.** Nb6 cannot bind RBD2 in open Spike*, as this would sterically clash with RBD3. **C,** Model of Spike*:Nb11 complex in the closed state. The minimal distance between adjacent Nb6 N- and C-termini is 71 Å (dashed line). **D,** Model of Spike*:Nb11 complex in the open state. The minimal distance between adjacent Nb6 N- and C-termini is 68 Å between Nb11 bound to RBD2 in the down-state and RBD3 in the up-state. For B, the model of Nb6 from A was docked into the cryo-EM map to enable modeling of distance between N- and C-termini. For C and D, a generic nanobody was docked into cryo-EM maps to model the distance between N- and C-termini.

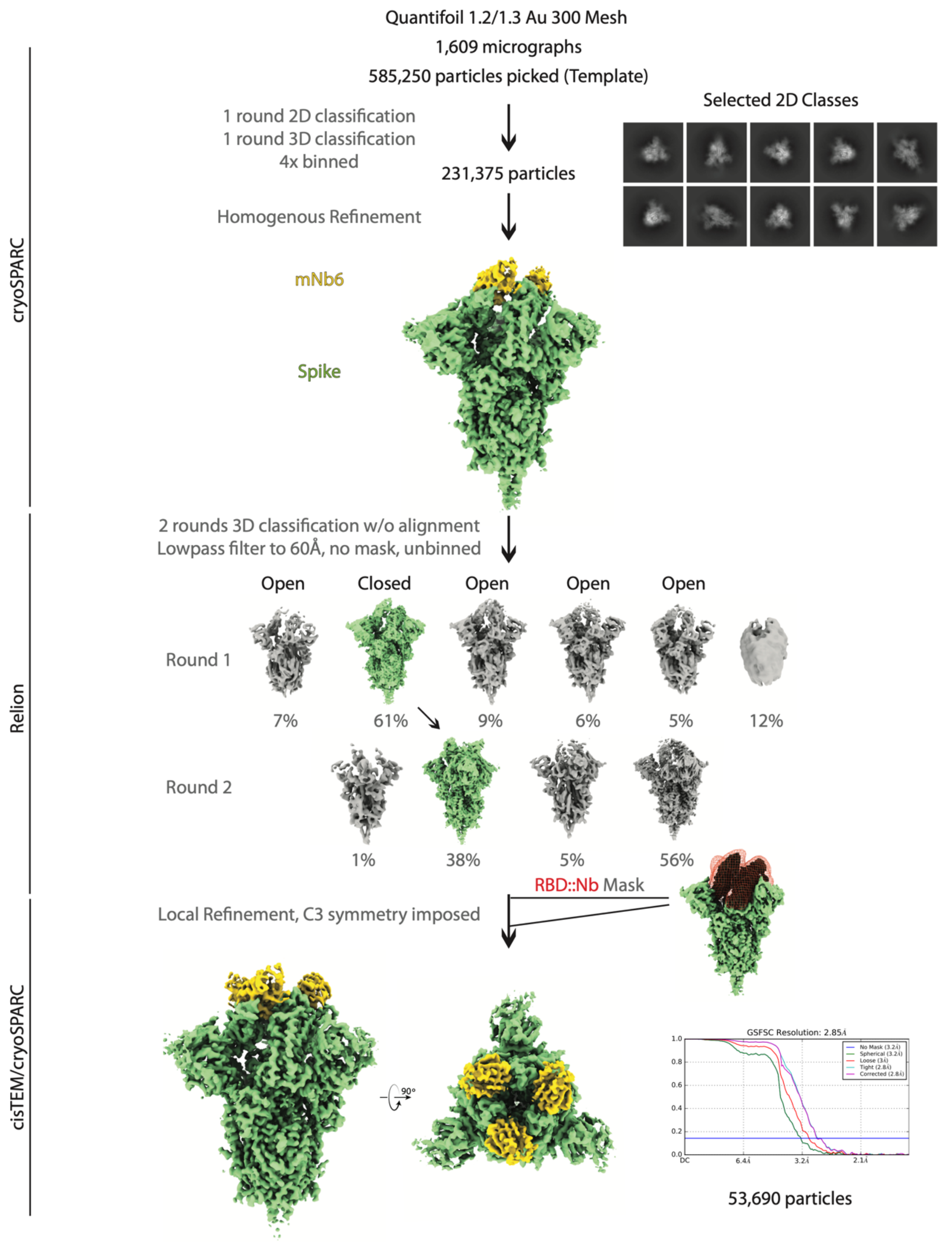

**Supplementary Fig. 7. CryoEM workflow for mNb6**

A flowchart representation of the classification workflow for the Spike*-mNb6 complex yielding a closed Spike* conformation. From top to bottom, particles were template picked from two separate collections with a set of 20Å low-pass filtered 2D backprojections of apo-Spike* in the closed conformation. Extracted particles were Fourier cropped to 96 pixels prior to 2D classification. Particles in Spike* 2D classes were selected for a round of heterogeneous refinement in cryoSPARC using a 20 Å low-pass filtered volume of apo-Spike* in the closed conformation and additional naïve classes for removal of non-Spike* particles. In RELION, particles in the Spike* 3D class were subject to two rounds of 3D classification without alignment into 6 classes using the same input volume from cryoSPARC 3D classification, low pass filtered to 60 Å, T = 8. Unbinned particles in the Spike*-closed conformation were exported into cisTEM for automatic refinement, followed by local refinement using a mask around the ACE2 RBD::Nanobody interface. Half maps from refinement were imported into cryoSPARC for local resolution estimation as shown in Supplementary Fig. 4.

#### Supplementary Table 1. CryoEM datasets

| Sample: | **Spike*-Nb6** | | **Spike*-Nb11** | | **Spike*-mNb6** |
|---|---|---|---|---|---|
| Spike* conformation: | Open | Closed | Open | Closed | Closed |
| EMDB: | XXXX | XXXX | XXXX | XXXX | XXXX |
| PDB: | | XXXX | | | XXXX |
| **Data collection and processing** | | | | | |
| Microscope/Detector | Titan Krios/Gatan K3 with Gatan Bioquantum Energy Filter | | | | |
| Imaging software and collection | SerialEM, 3x3 image shift | | | | |
| Magnification | 105,000 | | | | |
| Voltage (kV) | 300 | | | | |
| Electron exposure (e–/Å$^2$) | 66 | | | | |
| Dose rate (e–/pix/sec) | 8 | | | | |
| Frame exposure (e–/Å$^2$) | 0.55 | | | | |
| Defocus range (μm) | -0.8 to -2.0 | | | | |
| Pixel size (Å) | 0.834 (physical) | | | | |
| Micrographs | 5,317 | | 4,103 | | 1,609 |
| **Reconstruction** | | | | | |
| Autopicked particles (template-based in cryosparc) | 2,033,067 | | 1,204,855 | | 585,250 |
| Particles in final refinement | 40,125 (cisTEM) | 58,493 (cisTEM) | 21,570 (cisTEM) | 27,611 (RELION) | 53,690 (cisTEM) |
| Symmetry imposed | C1 | C3 | C1 | C1 | C3 |
| Map sharpening *B* factor (Å$^2$) | | -90 | | | -140 |
| Map resolution, global FSC (Å) | | | | | |
| FSC 0.5, unmasked/masked | 7.8/4.6 | 4.1/3.4 | 7.0/4.4 | 7.6/5.3 | 3.9/3.3 |
| FSC 0.143, unmasked/masked | 4.7/3.8 | 3.5/3.0 | 4.3/3.7 | 5.1/4.2 | 3.2/2.9 |
| **Refinement** | | | | | |
| Initial model used (PDB code) | | 6VXX, 3P0G | | | 6VXX, 3P0G |
| Model resolution (Å) | | | | | |
| FSC 0.5, unmasked/masked | | 3.5/3.1 | | | 3.2/2.9 |
| Model composition | | | | | |
| Non-hydrogen atoms | | 26904 | | | 27015 |
| Protein residues | | 3360 | | | 3360 |
| *B* factors (Å$^2$) | | | | | |
| Protein | | 97.0 | | | 57.5 |
| Ligand | | 107.4 | | | 85.7 |
| R.m.s. deviations | | | | | |
| Bond lengths (Å) | | 0.014 | | | 0.007 |
| Bond angles (°) | | 1.379 | | | 1.027 |
| Validation | | | | | |
| MolProbity score | | 1.99 | | | 1.71 |
| Clashscore | | 12.70 | | | 6.46 |
| Poor rotamers (%) | | 0.45 | | | 0.41 |
| EMRinger score | | 2.98 | | | 4.01 |
| CaBLAM score | | 3.11 | | | 2.95 |
| Ramachandran plot | | | | | |
| Favored (%) | | 94.49 | | | 94.92 |
| Allowed (%) | | 5.51 | | | 5.08 |
| Disallowed (%) | | 0 | | | 0 |

#### Supplementary Table 2. X-ray data collection and refinement statistics

| | mNb6 (PDB XXXX) |
|---|---|
| **Data collection** | |
| Space group | $P2_1$ |
| Cell dimensions | |
| *a*, *b*, *c* (Å) | 44.56, 71.25, 46.43 |
| α, β, γ (°) | 90.0, 114.93, 90.0 |
| Molecules in asymmetric unit | 2 |
| Resolution (Å) | 71.25 - 2.05 (2.09 - 2.05)[a] |
| $R_{sym}$ or $R_{merge}$ | 0.13 (0.94)[b] |
| $I / \sigma I$ | 7.2 (0.9) |
| Completeness (%) | 97.8 (96.6) |
| Redundancy | 6.4 (5.7) |
| CC (1/2) (%) | 99.8 (64.4) |
| **Refinement** | |
| Resolution (Å) | 71.25 – 2.05 |
| No. reflections | 104195 |
| $R_{work}$ / $R_{free}$ (%) | 21.16 / 24.75 |
| No. atoms | |
| Protein | 1798 |
| Ligand/ion | 21 |
| Water | 131 |
| *B*-factors | |
| Protein | 33.1 |
| Ligand/ion | 76.1 |
| Water | 42.2 |
| R.m.s. deviations | |
| Bond lengths (Å) | 0.07 |
| Bond angles (°) | 0.826 |
| Ramachandran plot | |
| Allowed (%) | 99.06 |
| Generous (%) | 0.94 |
| Disallowed (%) | 0 |

[a] Values in parentheses correspond to the highest resolution shell.
[b] $R_{merge} = \Sigma|I - \langle I \rangle|/\Sigma I$

#### Supplementary Table 3. Nanobody expression plasmids

| Plasmid | Nanobody | Plasmid backbone | Resistance Marker |
|---|---|---|---|
| pPW3544 | Nb2 | pet-26b(+) | kanamycin |
| pPW3545 | Nb3 | pet-26b(+) | kanamycin |
| pPW3546 | Nb6 | pet-26b(+) | kanamycin |
| pPW3547 | Nb8 | pet-26b(+) | kanamycin |
| pPW3548 | Nb11 | pet-26b(+) | kanamycin |
| pPW3549 | Nb12 | pet-26b(+) | kanamycin |
| pPW3550 | Nb15 | pet-26b(+) | kanamycin |
| pPW3551 | Nb16 | pet-26b(+) | kanamycin |
| pPW3552 | Nb17 | pet-26b(+) | kanamycin |
| pPW3553 | Nb18 | pet-26b(+) | kanamycin |
| pPW3554 | Nb19 | pet-26b(+) | kanamycin |
| pPW3555 | Nb24 | pet-26b(+) | kanamycin |
| pPW3557 | Trivalent Nb6, 20AA length GS linker | pet-26b(+) | kanamycin |
| pPW3558 | Trivalent Nb3, 15AA length GS linker | pet-26b(+) | kanamycin |
| pPW3559 | Trivalent Nb11, 15AA length GS linker | pet-26b(+) | kanamycin |
| pPW3560 | Bivalent Nb3, 15AA length GS linker | pet-26b(+) | kanamycin |
| pPW3561 | Bivalent Nb6, 15AA length GS linker | pet-26b(+) | kanamycin |
| pPW3563 | Trivalent mNb6, 20AA length GS linker | pet-26b(+) | kanamycin |
| pPW3564 | mNb6 | pet-26b(+) | kanamycin |

**QCRG STRUCTURAL BIOLOGY CONSORTIUM AUTHORS**

In addition to those listed explicitly in the author contributions, the structural biology portion of this work was performed by the QCRG (Quantitative Biosciences Institute Coronavirus Research Group) Structural Biology Consortium. Listed below are the contributing members of the consortium listed by teams in order of team relevance to the published work. Within each team the team leads are italicized (responsible for organization of each team, and for the experimental design utilized within each team), then the rest of team members are listed alphabetically. *CryoEM grid freezing/collection team**:** Caleigh M. Azumaya, Cristina Puchades, Ming Sun*, Julian R. Braxton, Axel F. Brilot, Meghna Gupta, Fei Li, Kyle E. Lopez, Arthur Melo, Gregory E. Merz, Frank Moss, Joana Paulino, Thomas H. Pospiech, Jr., Sergei Pourmal, Alexandrea N. Rizo, Amber M. Smith, Paul V. Thomas, Feng Wang, Zanlin Yu. *CryoEM data processing team: Miles Sasha Dickinson, Henry C. Nguyen*, Daniel Asarnow, Julian R. Braxton, Melody G. Campbell, Cynthia M. Chio, Un Seng Chio, Devan Diwanji, Bryan Faust, Meghna Gupta, Nick Hoppe, Mingliang Jin, Fei Li, Junrui Li, Yanxin Liu, Gregory E. Merz, Joana Paulino, Thomas H. Pospiech, Jr., Sergei Pourmal, Smriti Sangwan, Tsz Kin Martin Tsui, Raphael Trenker, Donovan Trinidad, Eric Tse, Kaihua Zhang, Fengbo Zhou. *Crystallography team: Nadia Herrera, Huong T. Kratochvil, Ursula Schulze-Gahmen, Michael C. Thompson, Iris D. Young*, Justin Biel, Ishan Deshpande, Xi Liu. *Mammalian cell expression team: Christian Bache Billesbølle, Melody G. Campbell, Devan Diwanji, Carlos Nowotny, Amber M. Smith, Jianhua Zhao*, Caleigh M. Azumaya, Alisa Bowen, Nick Hoppe, Yen-Li Li, Phuong Nguyen, Cristina Puchades, Mali Safari, Smriti Sangwan, Kaitlin Schaefer, Raphael Trenker, Tsz Kin Martin Tsui, Natalie Whitis. *Protein purification team: Daniel Asarnow, Michelle Moritz, Tristan W. Owens, Sergei Pourmal*, Caleigh M. Azumaya, Cynthia M. Chio, Amy Diallo, Bryan Faust, Meghna Gupta, Kate Kim, Joana Paulino, Jessica K. Peters, Kaitlin Schaefer, Tsz Kin Martin Tsui. *Bacterial expression team: Amy Diallo, Meghna Gupta*, *Erron W. Titus*, Jenny Chen, Loan Doan, Sebastian Flores, Mingliang Jin, Huong T. Kratochvil, Victor L. Lam, Yang Li, Megan Lo, Gregory E. Merz, Joana Paulino, Aye C. Thwin, Stephanie Wankowicz, Zanlin Yu, Yang Zhang, Fengbo Zhou. *Infrastructure team:* David Bulkley, Arceli Joves, Almarie Joves, Liam McKay, Mariano Tabios, Eric Tse. *Leadership team: Oren S Rosenberg, Kliment A Verba*, David A Agard, Yifan Cheng, James S Fraser, Adam Frost, Natalia Jura, Tanja Kortemme, Nevan J Krogan, Aashish Manglik, Daniel R. Southworth, Robert M Stroud. The QCRG Structural Biology Consortium has received support from: Quantitative Biosciences Institute, Defense Advanced Research Projects Agency HR0011-19-2-0020 (to D.A.Agard and K.A.Verba; B. Shoichet PI), FastGrants COVID19 grant (K.A.Verba PI),

Laboratory For Genomics Research (O.S.Rosenberg PI) and Laboratory for Genomics Research LGR-ERA (R.M.Stroud PI). R.M.Stroud is supported by NIH grants AI 50476, GM24485.
